# Impact of fluctuating environments on the fitness and robustness of evolving laboratory and industrial *Saccharomyces cerevisiae* strains

**DOI:** 10.64898/2025.12.05.692621

**Authors:** Cecilia Trivellin, Diana Ekman, Karl Persson, Misha Gupta, Lisbeth Olsson, Michael M. Desai

**Affiliations:** Department of Life Sciences, Division of Industrial Biotechnology, Chalmers University of Technology, Gothenburg, 412 96, Sweden; Department of Biochemistry and Biophysics, National Bioinformatics Infrastructure Sweden, Science for Life Laboratory, Stockholm University, Solna, Sweden; Department of Organismic and Evolutionary Biology, Harvard University, Cambridge, MA, 02138, United States

**Keywords:** Adaptive laboratory evolution, trade-offs, robustness quantification, pleiotropy, genome sequencing

## Abstract

Adaptation occurs through the selection of beneficial mutations enhancing fitness in a specific environment. However, since environments vary across time and space, mutations that are positively selected in one environment may be less beneficial or detrimental in others. Here, we investigate the evolution of microbial robustness (i.e. a consistent fitness across many diverse environments) through the adaptive evolution of two genetically distinct *Saccharomyces cerevisiae* populations in fluctuating conditions, followed by fitness assays and whole-genome sequencing. Our results indicate that the haploid laboratory strain S288C achieved higher average fitness than its parental strain, particularly when evolved in fluctuating environments compared to constant environments, but did not show increased robustness. In contrast, populations of the industrial diploid strain Ethanol Red failed to achieve significant fitness improvement under both fluctuating and constant evolution regimes but became more robust. Populations that adapted to fluctuating conditions acquired mutations in genes involved with cell morphology and protein degradation. Overall, our results emphasise the importance of parental traits in shaping fitness and robustness during adaptive laboratory evolution.

## Introduction

Adaptive laboratory evolution (ALE) has been employed over the years to improve strain fitness and to investigate biological phenomena (Dragosits & Mattanovich, 2013; Mavrommati *et al*, 2022; Sandberg *et al*, 2019). For example, integrating ALE with reverse genetic engineering has proven valuable in enhancing microbial tolerance to inhibitors found in lignocellulosic biomass (Menegon *et al*, 2022), and ALE performed in different cultivation environments revealed pleiotropic mechanisms in budding yeast (Chen *et al*, 2023). However, one commonly reported drawback is that adaptation in one environment can lower fitness in others (Kvitek & Sherlock, 2013; Wenger *et al*, 2011). This tradeoff occurs when beneficial mutations that are selected in one environment do not necessarily confer the same benefit, and may even be harmful, in a different environment.

To address fitness trade-offs in different environments, the integration of microbial robustness, a system’s ability to maintain consistent performance in different conditions (Olsson *et al*, 2022), is crucial. Microbial robustness has been studied from a phenotypic point of view (Trivellin *et al*, 2022; Torello Pianale *et al*, 2023), but its molecular mechanisms remain poorly understood. It is unclear whether robustness is regulated by individual genes or specific pathways, or if it results from more complex mechanisms, such as a modular metabolic architecture with feedback loops and redundant pathways (Kitano, 2004). Previous studies have pointed to heat shock proteins, sulfur metabolism and ribosomal proteins as potential robustness markers (Trivellin *et al*, 2024b; Félix & Barkoulas, 2015; Sangster *et al*, 2004; Levy & Siegal, 2008).

Evolution in fluctuating environments typically entails propagation over multiple generations in changing or oscillating conditions (Abdul-Rahman *et al*, 2021; Cooper & Lenski, 2010). The frequency of environmental fluctuations influences the evolution outcome. Evolution in constant environments generally favors specialists (population showing high fitness in one condition), while low-frequency fluctuations can encourage diverse or generalist offspring (populations with lower fitness than specialists but adaptable to diverse conditions). Higher frequency fluctuations require generalist phenotypes to avoid low fitness and mortality (Haaland *et al*, 2020). Previous studies have highlighted three aspects when evolving strains in fluctuating conditions: (i) evolution in chemostats has shown that fluctuating environments tend to favor genotypes with neutral fitness effects and lack extreme high- or low-fitness types, which helps preserve genetic diversity (Abdul-Rahman *et al*, 2021); (ii) deletion of genes that encode components of the ERAD pathways (targeting misfolded proteins for degradation) has resulted in high fitness in fluctuating conditions, probably due to the reduced rate of protein degradation (Abdul-Rahman *et al*, 2021); (iii) fluctuating environments can hinder selection’s efficiency across various parameters, complicating the distinction between beneficial and deleterious mutations (Cvijović *et al*, 2015). Furthermore, it has been shown that the specific history of encountered conditions influences the outcome of evolution, so fluctuating conditions with different histories might result in different evolutionary outcomes (Gonzalez & Bell, 2013).

Here, we set out to study the evolution of robustness in fluctuating environments, in order to address two objectives. First, we wanted to mitigate fitness tradeoffs across different environments (achieve higher robustness) by evolving strains under fluctuating conditions. Second, we aimed to describe the molecular mechanisms related to evolution in fluctuating environments and robustness by analyzing genomic variants in the evolved populations.

To achieve our objectives, we established an evolution experiment using two *Saccharomyces cerevisiae* strains that we propagated for 312 generations in 96-well plates in both constant and fluctuating environments that vary between fifteen different conditions. We selected two different strain backgrounds, one laboratory strain (S288C) and one industrial strain (Ethanol Red), to assess the impact of varying ploidy levels, genetic backgrounds, and the inherent fitness and robustness of the parental strains on the outcomes of experimental evolution. To evaluate the impact of fluctuation frequencies, we used three evolution regimes: one constant and two fluctuating (changing media every 8 or 80 generations). Finally, the specific fluctuation schedule was also changed within each regime. We assessed the fitness of the final samples of the evolution in 20 environments using a high-throughput phenotypic assay (Zackrisson *et al*, 2016). We also calculated robustness from the fitness data of the evolved samples using a robustness quantification method we recently developed (Trivellin *et al*, 2022). Whole genome sequencing was performed on all the evolved populations.

Our study combines adaptive laboratory evolution in fluctuating environments with robustness quantification to investigate fitness trade-offs, robustness, pleiotropy and molecular mechanisms of evolution in fluctuating conditions. We found that haploid and diploid strains evolved fitness and robustness in contrasting ways. Haploids with less fit parents significantly increased in fitness, while the more-fit parental diploid strain Ethanol Red instead enhanced its robustness over time. In contrast with previous findings in yeast, evolution in fluctuating conditions improved fitness more than constant conditions in certain measurement environments. We also identified cellular morphology and protein degradation as important mechanisms related to evolution in high-frequency fluctuating environments.

## Results

In this study, we wanted to address the effects of parental genotype and different types of fluctuating environmental conditions on adaptation of budding yeast populations. To do so, we initiated a total of 504 populations: 252 of one haploid *S. cerevisiae* strain, S288C (a standard laboratory strain), and 252 with a diploid strain, Ethanol Red (an industrial strain used for ethanol bioproduction). We evolved each of these parental strains using a standard batch culture protocol in 96-well microplates with daily 1:256 dilutions (Fig. 1A). Every 7 days, samples from each population were collected for future analysis.

**Figure 1.**
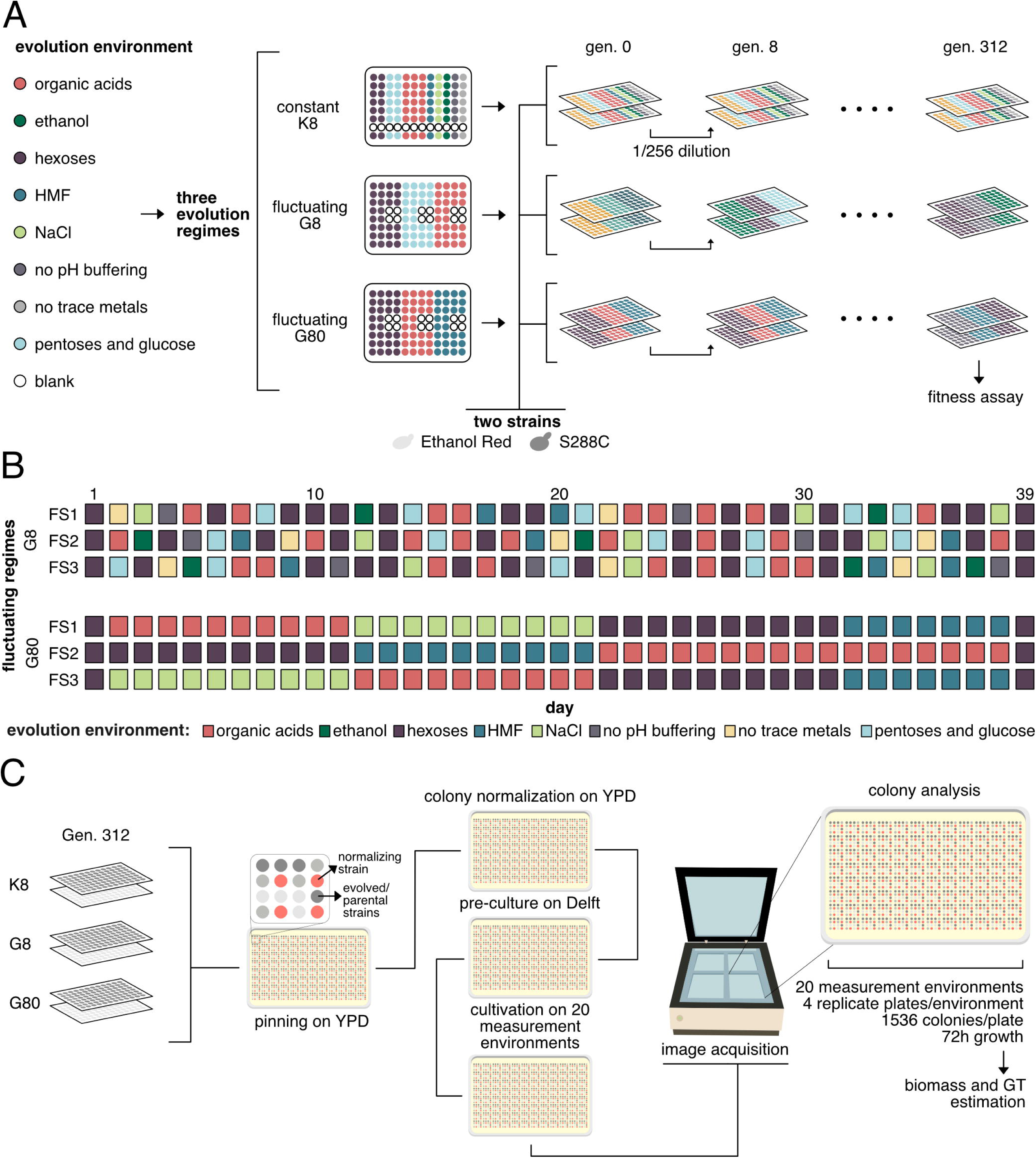
Schematic of experimental evolution and fitness assays. **A** Evolution setup: Two *Saccharomyces cerevisiae* strains were evolved in 96 well plates in three different regimes: in constant environments (K8) the strains were transferred in the same media daily; in fluctuating environments (G8) the strains were transferred every day in new media; in fluctuating environments (G80) the strains were transferred in new media every 8 days. Strains were transferred daily according to the dilution protocol showed at the top. The media used is showed on the left (Table 1, Materials and Methods). Each plate had a total of 12 blank wells (shown in white). **B** The fluctuation schedule is shown in each row; colors represent the different media (Table 1, Materials and Methods). Days correspond to columns. FS: fluctuation schedule. **C** Fitness assays (scan-o-matic): the end-samples from the evolution experiment were used to assess fitness. The six plates containing evolved strains were pinned on YPD in a 1536 colonies format according to a specific scheme (showed in the 4X4 squares). Red colonies represent colonies used for normalization of fitness across the plate. Colonies pinned on YPD were then pinned on YPD, then pre-cultured in Delft 2% glucose and then pinned in 20 different media (4X replicates) containing different chemicals (Table 2, Materials and Methods). Colony growth was monitored in scanners for 72 hours and the fitness was then assessed using standard normalization techniques (see Materials and methods for details).

**Table 1.** List of chemicals and concentrations added to the evolution media.

| **Chemical** | **Concentration (g/L)** |
| --- | --- |
| D-(+)-Glucose | 65, 20, 5 |
| D-(+)-Xylose | 8 |
| L-(+)-Arabinose | 4 |
| D-(+)-Mannose | 12 |
| Formic acid | 2 |
| Acetic acid | 2 |
| DL-Lactic acid | 7 |
| Levulinic acid | 4 |
| 5-Hydroxymethylfurfural | 1 |
| Ethanol | 20 |
| NaCl | 25 |
| No Trace Metals |  |
| No pH buffer |  |

*D-(+)-glucose 20 g/L was used in all the media as carbon source except in presence of D-(+)-xylose and L-(+)-arabinose in which the concentration of glucose was reduced to 5 g/L and no glucose was present when using D-(+)-mannose.*

**Table 2:**
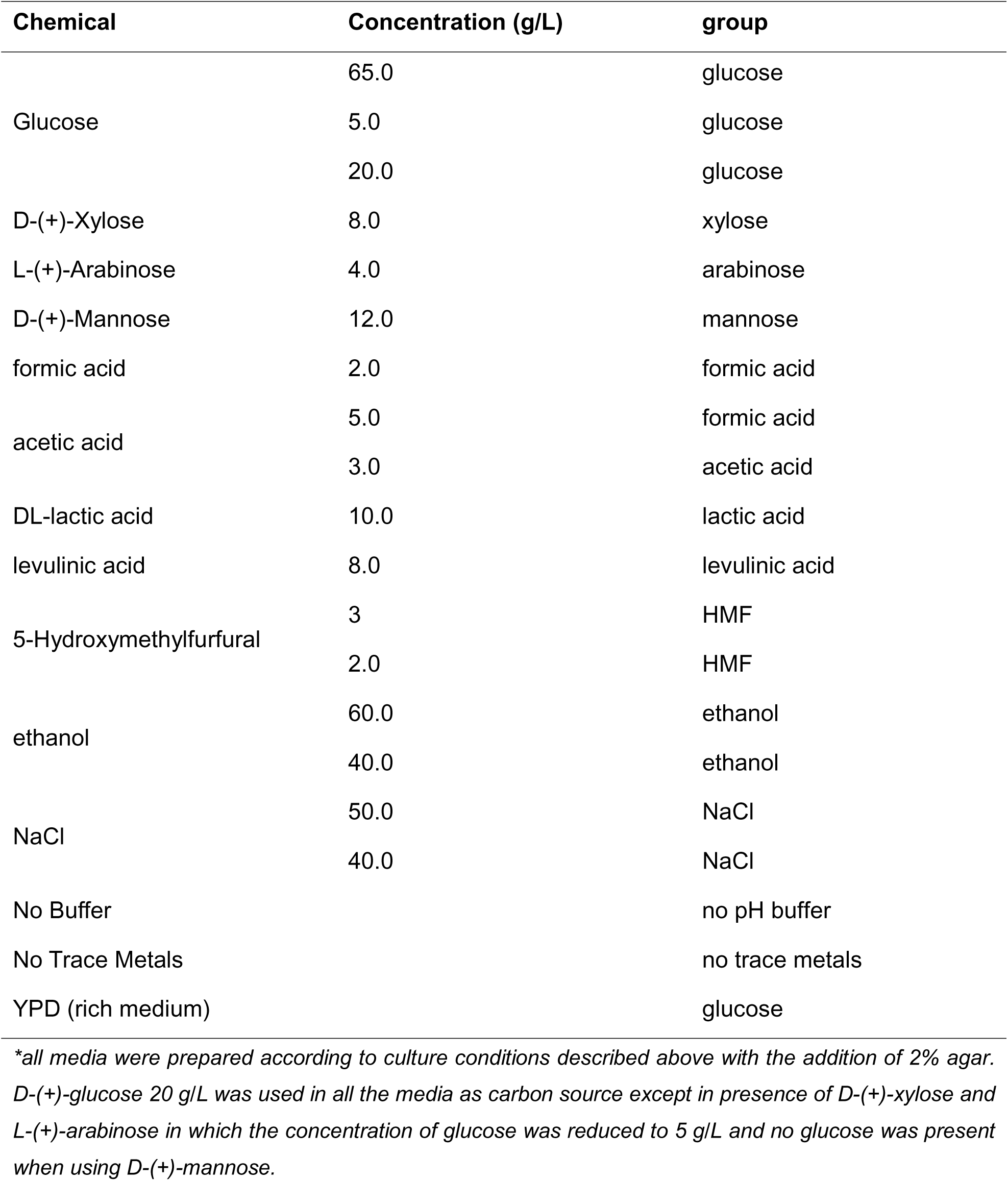
Media composition for phenotypic assays.

We implemented three distinct evolution protocols, which we refer to as K8, G8, and G80. For the K8 protocol, we propagated 7 replicate lines for each of the two parental strains in each of 12 different constant environments (one of 12 different media conditions). For G8, we propagated 28 replicate lines for each of the two parental strains in each of 3 different environmental regimes that involve a different media condition each day (every 8 generations). For G80, we propagated 28 replicate lines for each of the two parental strains in each of 3 different environmental regimes that involve a different media condition every 10 days (every 80 generations). Details of these propagation regimes are illustrated in Fig. 1B.

After 312 generations of evolution, we sequenced whole-population (metagenomic) samples from all 504 populations, along with two clones isolated from each of 96 evolved populations (for each parental strain, 2 evolved lines from each of the 12 K8 environments and four evolved lines from each of the three G8 and three G80 regimes). We implemented a bioinformatics pipeline to identify mutations arising in each of these whole-population and clonal samples (for details see Methods). This sequencing data revealed that, likely early in the experiment, 274 of the populations became contaminated. We excluded these lines from all further analysis, leaving a total of 230 evolved lines.

We analyzed changes in growth phenotypes in each evolved line across a range of 20 media conditions using a scan-o-matic high-throughput phenotyping setup (Zackrisson *et al*, 2016), which uses imaging to estimate the maximum total biomass and generation time in each colony through 72 hours of growth (Fig. 1C). Specifically, we first pinned whole-populations samples and single colonies isolates from generation 312 onto YPD plates in a 1536 colony format. We pinned a control strain, CEN.PK113-7D, across the plate for spatial normalization, while parental strains were positioned differently for spatial comparison of fitness. We then pinned on YPD for strain recovery and Delft medium for preculturing, before pinning onto plates containing 20 different media conditions (see Methods). We monitored colony growth for 72 hours using scanners (four replicate plates in each scanner) and estimated the spatially normalized maximum biomass and generation time of each colony using the scan-o-matic software (see Methods for details). We use the estimate of maximum biomass as a proxy for the fitness of the corresponding strain in that media condition. We note that the biomass represents growth of colonies on solid media, which does not perfectly reflect the selection pressures that may exist during growth in corresponding liquid media that was used during the evolution experiment.

Although we evaluated fitness in terms of biomass, we also measured the generation time of each evolved population in each measurement environment compared to the generation time of the parental strains (Appendix Figure S1). However, considering the modest variability of the different populations, both in different evolution replicates and in the growth assay replicates, we could not significantly distinguish between changes in population fitness due to adaptation from experimental noise and assay variation. For this reason, we decided to report results in terms of biomass. We also plotted the relative biomass vs relative generation time (Appendix Figure S2). We observed that, for the evolution in fluctuating regimes, there seems to be a trade-off between fitness gain in terms of biomass or generation time. Ethanol Red, in fact, shows a higher percentage of populations with increased generation time compared to S288C. The opposite can be said for S288C in terms of biomass. However, this trade-off was not supported by a significant inverse linear correlation and cannot be validated due to the variability discussed above.

### Adaptation to certain home environments led to overall fitness gains in away environments

Adaptation to each of the 12 different constant environments (K8) did not result in a significant increase in fitness in the same environment (home environment) (Fig. 2A), even considering the modest variability of replicate populations (Appendix Figure S3). Interestingly, the only evolution environments that resulted in a fitness increase in the ‘home’ environment also showed a fitness increase in ‘away’ environments (environments were strains did not evolve in). For example, for the industrial strain Ethanol Red, evolution in glucose 20 g/L impacted overall fitness positively, especially when the evolved samples were tested in acids. For the laboratory strain S288C, evolution in formic acid resulted in a fitness increase of at least 50% in all the other measurement environments. For S288C, evolution in lactic acid also caused an increase in fitness towards acid environments and in HMF while evolution with no pH buffering resulted in a fitness decrease in all measurement environments. Overall, when plotting the first two principal components of each population (fitness values in all environments) the adapted populations showed similar fitness patterns in all evolution environments, but those listed above (glucose for Ethanol Red and formic acid and no pH buffer for S288C) clustered away from the rest, indicating a unique behavior (Fig. 2B).

**Figure 2.**
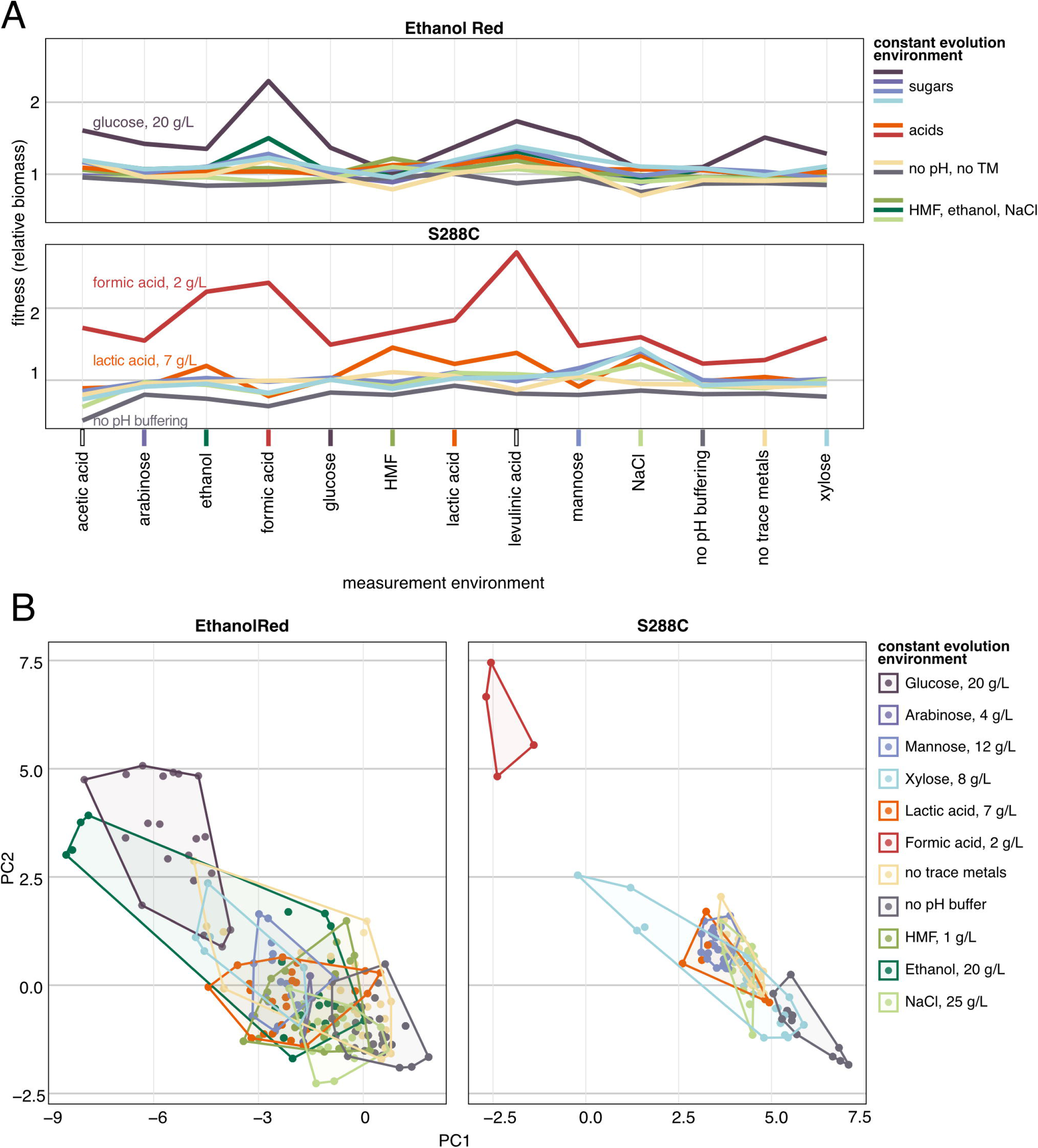
Fitness of strains evolved in constant environment (K8). **A** Each line connects the average fitness ratio (evolved strain biomass divided by corresponding parental strain biomass) across replicate populations measured in the same environment (x-axis). Each line color corresponds to an evolution environment that was kept constant throughout evolution. Environments in which fitness is significantly different from 1 are described by the text on the left of the plot. The x-axis groups measurement environments based on chemical similarity in the fitness assay (Table 2). The different colors correspond to the different media. No color means there was not a corresponding evolution environment. The two facets show results for the two different strains. **B** PCA was performed on fitness values of all the populations evolved in constant environments (each dot) across measurement environments. Populations that evolved in the same environment are clustered by a convex hull and colored differently. PCA was run separately for the two strains (facets). The axis represents the first two principal components.

### Laboratory strain showed a higher fitness gain following evolution in fluctuating versus constant environments

The industrial strain, Ethanol Red, did not show a significant fitness gain under the three evolution regimes (Fig. 3A). For the laboratory strain S288C, instead, the mean fitness gain across environments was significantly higher when the strains evolved in fluctuating regimes than constant regimes (Fig. 3A). This result reflects previous observations found when evolving bacteria in fluctuating temperatures (Ketola *et al*, 2013), but is contradictory to previous studies in yeast, where evolution in fluctuating environments increased fitness when the measurement environment itself was fluctuating, but negatively affected fitness in a constant measurement environment compared to strains evolved under constant conditions (Dhar *et al*, 2013). This outcome may be due to our experiment involving fluctuations across 15 different environments, in contrast to previous studies that used fewer than five media(Ketola *et al*, 2013; Dhar *et al*, 2013). The two distinct behaviors for Ethanol Red and S288C are also showed with the principal component analysis (run on the fitness values across environments) which resulted in two almost not overlapping and divergent clusters when highlighting the two strains (Fig. 3B).

**Figure 3.**
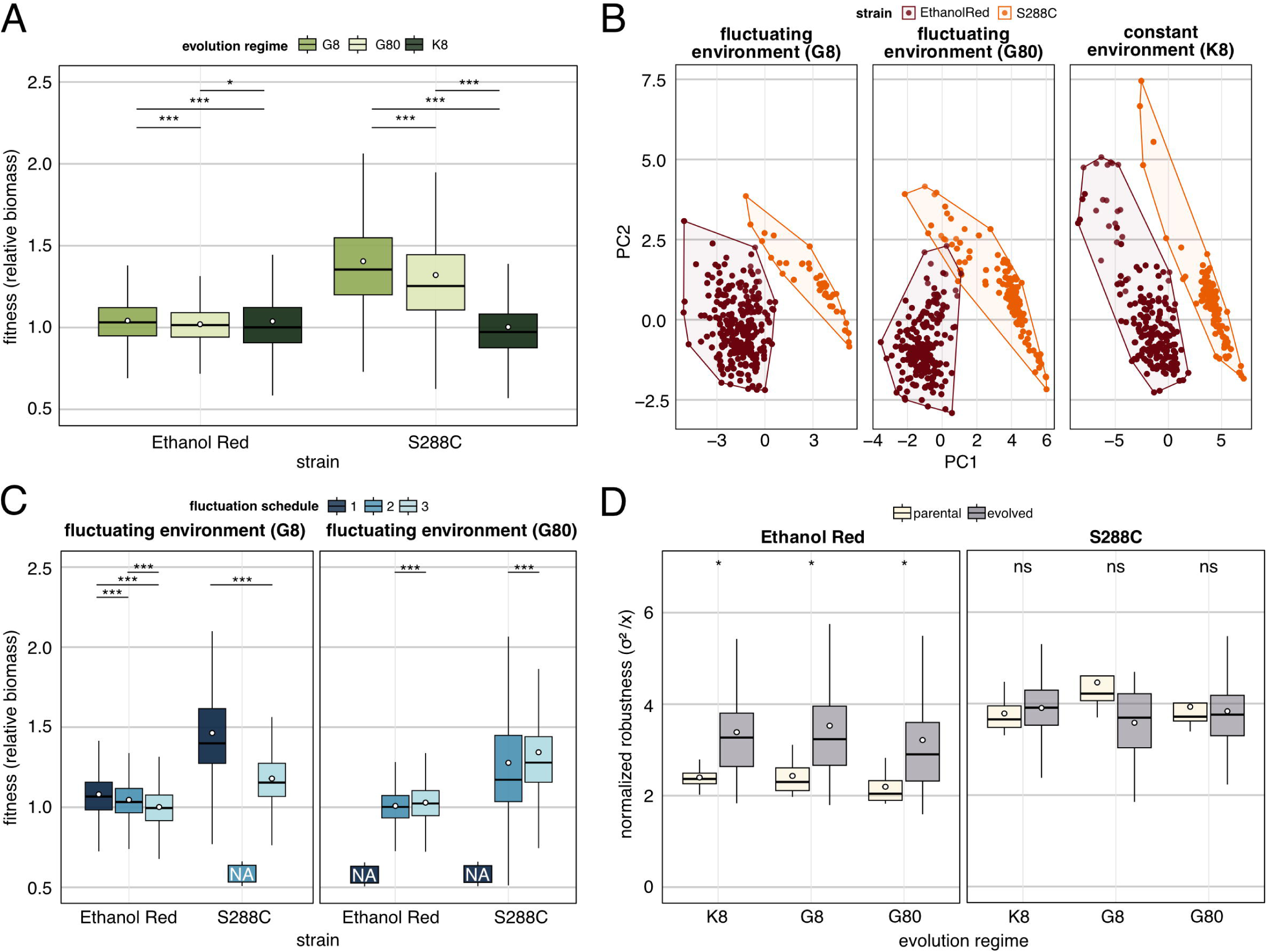
Fitness of evolved strains. **A** The y-axis depicts the relative fitness (evolved strain biomass divided by corresponding parental strain biomass). The x-axis represents the strains. Each boxplot shows the summary statistics for all the populations in one evolution regime (colors) across all measurement environments. The white dot shows the mean of the distribution. Significance stars represent p-values calculated with a Wilcox test (*** p-value <0.001; ** p-value <0.01; * p-value < 0.05; NS not significant). **B** PCA was performed on fitness values of all the evolved populations (each dot) across measurement environments. Strains are clustered by a convex hull and colored differently. PCA was run separately for the three evolution regimes (facets). The axis represents the first two principal components. **C** Similar to **A** but each boxplot shows the summary statistics for each fluctuation schedule (colors) divided by fluctuating evolution regime (facets). The NA boxplot shows missing values due to contamination (see Materials and Methods). **D** The y-axis shows the robustness calculated with Eq. 1 (see Materials and Methods) for each evolution regime (x-axis). Robustness was calculated for each evolved population and parental strain (the two boxplots of different colors) across the measurement environments. The white dot corresponds to the mean of the distribution. Significance stars are as in **A**.

The strain S288C showed a lower mean fitness across environments for the G80 than the G8 regime. However, when the adapted strains evolved in G80 and were then measured in the environments that were mainly present in that regime (HMF, NaCl, Fig. 1B) the mean fitness was higher than those evolving in the G8 regime (Appendix Figure S4). This may indicate that beneficial mutations that emerged in those environments were likely fixed in the population and continued to confer an advantage even after the medium changed. However, this was only true for those two conditions and not for glucose 65 g/L and formic acid 2 g/L.

The fitness of the adapted strains was not only a result of the random fluctuations in environments: the fluctuation schedule (Fig. 3C) also had a notable impact. In the case of G8, fluctuation schedule 1 consistently yielded the highest fitness increase for both strains. A clear difference from the three lines in G8 was that in fluctuation schedule 1, strains were exposed for longer periods of time to hexoses. In the case of G80, fluctuation schedule 3 resulted in higher fitness for both strains. Surprisingly, fluctuation schedule 3 resulted in a greater increase in fitness when exposed to acids, despite fluctuation schedule 2 being subjected to acids for a longer duration (Fig. 1B). A more detailed overview of the fluctuation schedules and strains in all environments is available in Appendix Figure S5.

### Fitness gain is influenced by parental fitness

We analyzed the parental fitness of both industrial and laboratory strains (Appendix Figure S6). We found that Ethanol Red showed an overall higher fitness (biomass) than S288C, especially in high concentrations of glucose and stressors (Appendix Figure S6A). This is consistent with our expectation that industrial strains, such as Ethanol Red, have higher fitness in presence of stressors such as acids, aldehydes, and ethanol (Kang *et al*, 2019). The generation time of the parental strains was similar in most measurement environments, except for HMF where S288C was close to 10h compared to a mean of 3h for Ethanol Red (Appendix Figure S6B). Previous phylogenetic studies confirmed that Ethanol Red is closely related to S288C (Gronchi *et al*, 2022). We performed whole-population genome sequencing analysis and confirmed that Ethanol Red is diploid, whereas S288C is haploid. When mapped against the S288C reference genome, Ethanol Red showed 79,863 variants, consistent with prior observations (Wallace-Salinas *et al*, 2015) (Appendix Figure S7 right). The majority of variants were single-nucleotide polymorphisms. Major deletions in the telomeric regions were also identified (Appendix Figure S7 left, Appendix Table S1). As previous studies have shown that the strain genetic background could influence the outcome of evolution (Chandler *et al*, 2014; Goldstein & Ehrenreich, 2021), we investigated whether parental strain fitness was negatively correlated with the fitness increase after 300 generations of evolution. We found that when strains evolved in constant environments, the correlation was insignificant (Appendix Figure S8). In contrast, for G8 and G80 the relative fitness (biomass) was higher when parental strains were less fit, resulting in a significant negative correlation, with spearman coefficient lower than −0.6 (Appendix Figure S8). The correlations are affected by the measurement environments we chose for fitness assays.

### Industrial strain Ethanol Red increased robustness after evolution

To test our hypothesis that adaptation to fluctuating regimes would result in higher robustness, we used the data from the fitness measurements (biomass) to calculate our index of strain robustness (Eq. 1, Methods, modified from (Trivellin *et al*, 2022)). The robustness was calculated across 20 measurement environments, using fitness data (biomass) of evolved and parental strains separately.

The broad fitness distribution we observed among the evolved strains, likely driven by divergent evolution in replicate populations, was reflected in the similarly broad distribution of their robustness (Fig. 3D). The robustness is showed separately for parental and evolved strains to simplify interpretation. The S288C parental strain showed a higher robustness compared to Ethanol Red, possibly due to consistently low fitness across all measurement environments (Appendix Figure S6A). Evolution did not result in increased robustness of S288C, no matter the regime, but we found it did increase robustness in Ethanol Red (despite variability among populations), nearly reaching the level of the S288C parental strain (Fig. 3D). Variation in robustness was more pronounced among strains than across evolution regimes, suggesting that that robustness is influenced primarily by the parental strains rather than the evolution regime.

### Variants and functional enrichment analysis reveals that adaptation to fluctuating environments is associated with morphology changes and protein degradation

We analyzed genomic variants from each of 192 single clones, using a bioinformatic pipeline (as described in the Methods section). The sequenced samples exhibited a higher proportion of non-synonymous variants than synonymous ones (Fig. 4A). Mutations resulting in stop codons appeared in clones evolving in K8 and G8 but not in G80. In general, evolution in constant environments showed a higher variant count than G8 and G80, and S288C showed a higher variant count in G80 than Ethanol Red (Fig. 4B).

**Figure 4.**
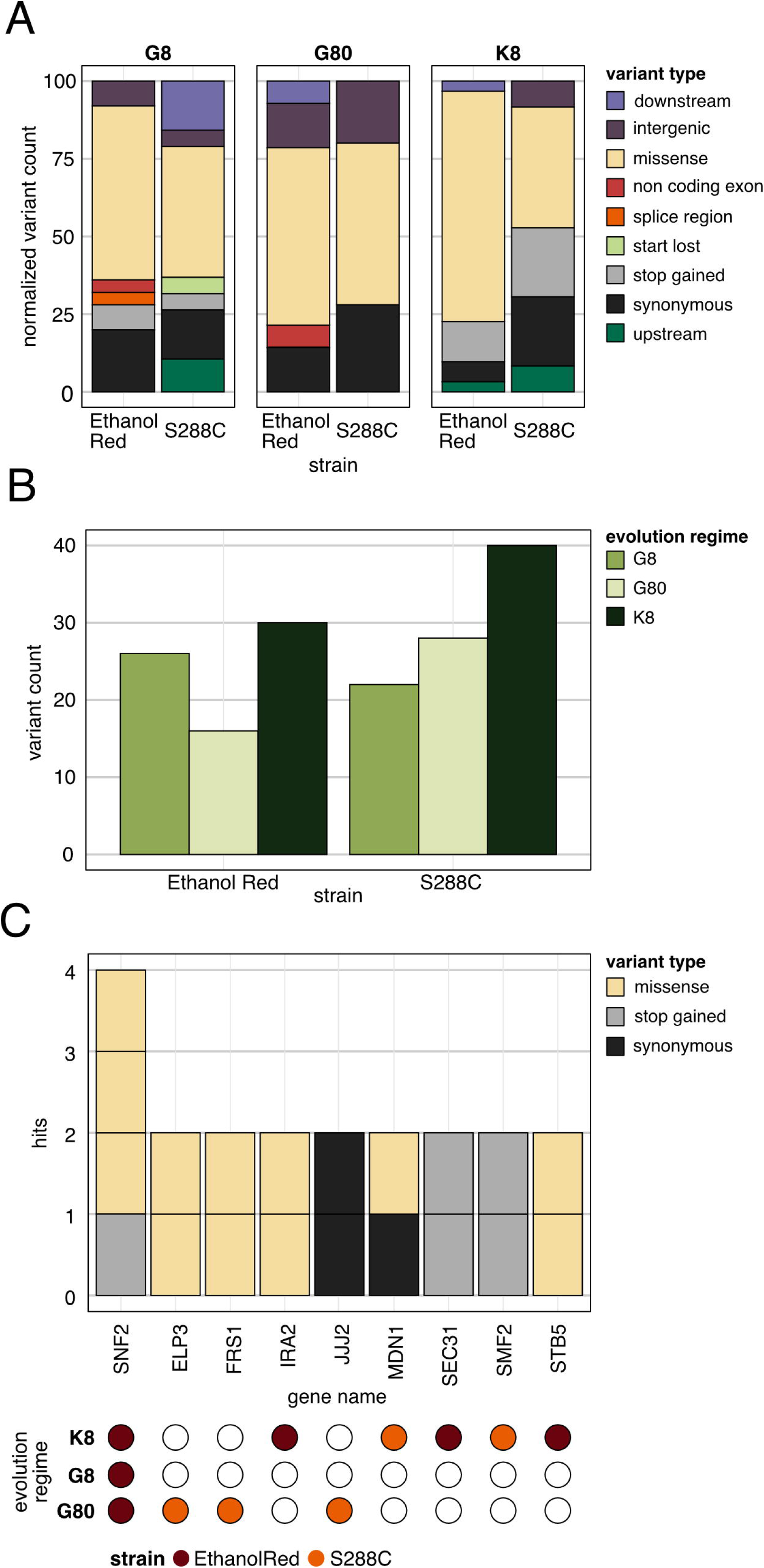
Sequence variant analysis. **A** Breakdown of types of variants of the end-evolution samples (single-colonies). Strains are shown on the x-axis and facets correspond to evolution regimes. Variant types are shown in different colors. The variant count per strain and evolution regime was normalized with the total count of variants (total=100). **B** Total count of variants for each strain (x-axis) and evolution regime (color). **C** Multi-hit genes: the x-axis represents genes that mutated in more than one sample. Bars were colored based on the variant type. The bottom part of the graph represents the evolution regime (rows) and strain background (color) where the mutations arose.

To test whether evolution within the same regime followed parallel trajectories, we identified genes with recurrent mutations across samples, classifying those with more than two counts as "multi-hit genes". Out of 152 unique variants, only 9 genes were affected more than once across different samples (Fig. 4C). Multi-hit genes were generally found in clones from the same evolutionary regime, with the sole exception of *SNF2*, which was mutated across all three evolutionary experiments. No gene hits were shared between the two strains. Notably, *SNF2* was also the only multi-hit gene detected in the G8 environment. Most mutations in multi-hit genes were non-synonymous. We examined protein-protein interactions of multi-hit genes using the STRING database (Szklarczyk *et al*, 2023), which incorporates both physical interactions and functional associations. Multi-hit genes were categorized by evolutionary regime and strain. The analysis revealed no significant interactions among the proteins. Examining structural variants revealed two distinct samples of the S288C evolving in G80 with significant deletions spanning multiple genes: HAL5, TPK1, YJL163C and JJJ2. These deletions were positioned on chromosome X (6525 bp). The deleted genes were mainly associated with proteins of unknown function, but the deletion in S288C was linked to genes associated with nutrient response (for example HAL5). Despite rigorous analysis of previous null mutant studies, no clear phenotypic benefits from these deletions were identified.

Next, to confirm if mutations that arose during evolution would fix more frequently than expected by chance, population hits and multiplicity values were calculated. We define a “hit” as a nonsynonymous mutation that becomes fixed by the final timepoint and multiplicity of a gene as the number of hits observed in that gene across all sequenced populations, normalized by its relative target size (Good *et al*, 2017; Johnson *et al*, 2021). The distribution of the population hits and multiplicity was tested against a multinomial null distribution simulating random hits to genes (Appendix Figure S9). A non-parametric test evaluating the differences between test and null distribution revealed a non-significant difference (p-value=0.36 in both cases). The non-significance could also be attributed the to the small number of multi-hit genes that we found in our experiment (9), compared to, for example, 42 found by (Johnson *et al*, 2021).

We subjected genes associated with genomic variants to functional enrichment analysis to provide a comprehensive overview of the principal cellular processes and components associated with the outlined mutations. We used the g:profiler tool (Kolberg *et al*, 2023), a collection of different ontology terms, and selected uniquely adjusted p-values corresponding to significant enrichment (p_adj_ < 0.05) (Appendix Figure S10 and Appendix Table S2). When analyzing genes that mutated in the different evolution regimes (no matter the strain), the G80 regimes were associated uniquely with tRNA uridine (34) acetyltransferase activity while both the GO terms associated with the G8 experiment were associated with regulation of cell adhesion in biofilm formation. The set of genes belonging to the K8 set did not result in significantly enriched terms. If the ‘strain’ variable was considered in the analysis of GO terms, the S288C strain showed uniquely glycan degradation in the G8 regime, while Ethanol Red mutated genes were associated with regulation of distinct processes in the K8 regime, and two transcription factors: Tec1p in G8 and Abf1p in G80. To summarize, evolution in fluctuating environments, especially at high frequency of fluctuations, targeted genes associated with change in morphology and the ERAD pathway (as previously observed in fluctuating regimes (Abdul-Rahman *et al*, 2021)).

## Discussion

The primary objective of this study was to explore how different strains of *Saccharomyces cerevisiae* evolve in both constant and fluctuating environments, with the goal of assessing the impact of fluctuating selective pressures on fitness and microbial robustness. We evolved 230 independent populations for approximately 312 generations. We then grew samples from the evolution endpoints in 20 diverse conditions, measured fitness and robustness, and sequenced their genomic DNA.

Earlier work has consistently observed that evolution in stable environments leads to enhanced fitness within the same environment (Dolpatcha *et al*, 2023; Gong *et al*, 2023; Sánchez-Adriá *et al*, 2023; Pereira *et al*, 2019), but sometimes leads to trade-offs resulting in lower fitness in different environments (Persson *et al*, 2022). In the present study, we did not observe a fitness trade-off in ‘home’ vs ‘away’ environments. Evolution in a single environment either improved fitness across the majority of measurement environments or had overall no effect. We also observed that evolution without pH buffering decreased fitness in all environments. The absence of an observed trade-off in fitness across environments may be because the conditions did not impose strong divergent selective pressures and thus did not favor the evolution of condition-specific mechanisms that could lead to trade-offs in ‘away’ environments. In contrast, we did observe an overall increase in fitness across all tested conditions, which could be explained by synergistic pleiotropy where mutations acquired by the populations positively affect fitness across multiple environments. Whether pleiotropy promotes robustness remains unclear. One could argue that pleiotropic effects might increase population robustness in the short term but could also make the population more vulnerable to deleterious mutations over longer evolutionary timescales.

The highest fitness gain was observed in populations of S288C evolved in fluctuating environments. Previous research in yeast has suggested that fluctuating environments lead to increased fitness only if the measurement environment also fluctuates (Dhar *et al*, 2013), while in our case, the fitness increase was present in all measurement environments. Like in the case of evolution in certain constant environments, evolution in fluctuating environments might have favored pleiotropy or could be attributed to the anticipatory behavior developed by frequently encountering different stressors (G8). It is also plausible that mutations that fixed in fluctuating environments are linked to a general stress response or a reaction to reactive oxidative species, which are common in the evolution conditions used in this study (Ndukwe *et al*, 2020; Ikner & Shiozaki, 2005; Pérez-Gallardo *et al*, 2013). Fluctuating conditions at a lower frequency (80 generations instead of 8) may give enough time for fixation of condition-specific beneficial mutations in the population, resulting in higher fitness in all the measurement environments that were encountered already in the evolution (in this case higher fitness towards HMF and NaCl enriched environments). However, changing the media every 8 generations might be too rapid for condition-specific beneficial mutations to fix, so the mutations observed after 300 generations could have reached appreciable frequencies partly through genetic drift rather than strong selection. Notably, certain fluctuation schedules consistently improved fitness across independent experiments, suggesting that some schedules may be inherently more beneficial. While we could not identify clear patterns underlying this effect, it warrants further investigation.

Fitness effects varied significantly among strains, with S288C exhibiting substantial fitness increase during evolution, while Ethanol Red fitness remained similar or even decreased in certain conditions, such as in NaCl enriched environment. This underscores the impact of parental traits on evolutionary outcomes, as highlighted by the negative correlations between parental fitness and fitness gain. Selective pressures act differently on the different strains, suggesting that the pressures benefiting S288C might not significantly impact Ethanol Red. In contrast to the lab strain, Ethanol Red is diploid. Previous studies have highlighted the impact of ploidy on adaptation, suggesting that diploid strains with mutations that elevate the spontaneous mutation rate can give rise to beneficial mutations that are inaccessible to haploid strains (Thompson *et al*, 2006). In our study, the chosen selective pressure may not have been sufficient for beneficial mutations to prevail in the Ethanol Red strain. Deleterious mutations might have accumulated, first masked by other factors such as higher copies of the same gene or phenotypic capacitors, and then released when strains were cultivated in the measurement environments in the phenotypic assays (for example Ethanol Red grown in NaCl). In Ethanol Red, the effect of beneficial mutations could have been offset by deleterious ones, resulting in no observable change in fitness in the final phenotypic assays.

Ethanol Red, which did not show increased fitness, exhibited increased robustness. The conditions used in the evolution seemed to level out fitness (sometime reducing fitness in some measurement conditions) across environments, enhancing overall strain robustness. This contrasted with S288C, where the robustness decreased. As a result of the present study, we observed that the robustness outcomes differed more across strain backgrounds than across evolution regimes. Since both fitness and robustness appear to be correlated with parental traits, this study raises the question of whether long-term evolution under the same selective pressures would lead to a gradual increase in overall fitness, as observed in Lenski’s *E. coli* experiment where fitness gains showed no signs of plateauing even after 50,000 generations (Wiser *et al*, 2013; Good *et al*, 2017), or whether the strains would instead become more robust, balancing fitness across different conditions -or possibly both.

Variant analysis and gene ontology analysis revealed that biological regulation, protein degradation, and morphology are the predominant processes relevant in the current evolution experiment, especially under fluctuating conditions. Protein degradation has been associated with adaptation to fluctuating environments (Abdul-Rahman *et al*, 2021), where a reduced degradation rate may confer an advantage. Proteins that are dispensable or harmful in one environment can become beneficial in another. Morphological adaptations such as biofilm formation, flocculation, and hyphal growth have previously been shown to provide advantages in stressful environments (Bouyx *et al*, 2021). These traits may be particularly beneficial under fluctuating conditions. For instance, flocculation can promote rapid resource sharing or dispersal when the environment changes, while hyphal growth enables exploration of new niches or access to nutrients that vary over time (Kim & Rose, 2015). By contrast, in stable environments such plasticity may offer little advantage and could even impose unnecessary energetic costs.

Multi-hit gene analysis revealed recurrent mutations in SNF2, ELP3, FRS1, and JJJ2 under the fluctuating regime. Several factors may explain why these genes mutated repeatedly. For example, Snf2p, as part of the SWI/SNF chromatin remodeling complex (Shivaswamy & Iyer, 2008), might enable rapid transcriptional reprogramming by modulating chromatin accessibility. Elp3p, a histone acetyltransferase subunit (Fernández-Vázquez *et al*, 2013), could enhance translation efficiency via tRNA modifications. Frs1p, a phenylalanine-tRNA ligase (Mohler *et al*, 2018), could contribute to survival in fluctuating conditions by fine-tuning the expression of stress-responsive genes. Jjj2p, a protein of unknown function, contains a conserved J domain characteristic of Hsp40 proteins, enabling interaction with Hsp70 to stimulate ATP hydrolysis, and may help maintain protein homeostasis by enhancing Hsp70 responsiveness during fluctuating environmental conditions (Walsh *et al*, 2004). Together, these genes could contribute to survive efficiently under dynamic and challenging conditions. However, these possibilities remain speculative. Reverse engineering of the identified mutations would help determine whether they indeed confer an advantage in fluctuating environments. Similarly, reconstructing mutations from clones that showed increased robustness could provide insights into the molecular mechanisms underlying microbial robustness.

In conclusion, our findings suggest that parental traits play a crucial role in shaping evolutionary outcomes, influencing both fitness and robustness. Our results indicate that robustness evolution may occur when the parental strain already exhibits high fitness, as observed for Ethanol Red. Ploidy may also contribute to the evolution of robustness, although additional strains are required to confirm this. Since our results stem from a short-term evolution experiment, it will be important to assess whether the same patterns hold in long-term experiments and under more diverse environmental conditions. Adaptation to fluctuating environments may arise through pleiotropic effects, morphological changes, or reduced protein degradation, as suggested by the variant analysis. Further experiments will be necessary to validate these mechanisms and potentially link them to robustness.

## Materials and Methods

### Strains

Two *Saccharomyces cerevisiae* strains were used for this study: one laboratory haploid strain, S288C (University of Milano Bicocca) with genotype MATα, SUC2, *gal2, mal, mel, flo1, flo8-1, hap1, ho, bio1, bio6* and one industrial strain, Ethanol Red, diploid and kindly provided by Société Industrielle Lesaffre, Division Leaf.

### Culture conditions

Fifteen different media were prepared using Delft minimal medium (Bruinenberg *et al*, 1983) as the base. The medium contained 5 g/L (NH_4_)_2_SO_4_, 3 g/L KH_2_PO_4_, 1 g/L MgSO_4_·7H_2_O, 1 mL (in 1 L solution) trace mineral solution (Appendix Table S3), and 1 mL (in 1 L solution) vitamin solution (Appendix Table S4). The medium was adjusted to pH 5 with potassium hydroxide and buffered with 100 mM potassium hydrogen phthalate. To simulate perturbations encountered during bioethanol production, the compounds listed in Table 1 were added to the base medium (Trivellin *et al*, 2024a).

### Evolution

Cultures were propagated in 128 µl of media in sterile round-bottom polypropylene 96-well plates (Greiner Bio-One 655261) with lids. Populations were grown at 30°C, 1300 rpm orbital shaking. A total of six 96-well plates were used in the evolution. Two plates (one strain for each plate) had 12 control conditions divided into columns with 7 biological replicates + 1 blank well (Delft with glucose 20 g/L, mannose 16 g/L, xylose 8 g/L, arabinose 4 g/L, formic acid 2 g/L, lactic acid 7 g/L, acetic acid 2 g/L, HMF 1 g/L, ethanol 20 g/L, sodium chloride 25 g/L, Delft with 20 g/L glucose without potassium hydrogen phthalate and Delft with 20 g/L glucose without trace metals). Two plates (one strain each) were divided into three evolution lines (different fluctuation schedules) each with 28 biological replicates and 4 blanks. The three fluctuation schedules corresponded to three different sequences by which media change at every transfer (every 8 generations). The remaining two plates (one strain each) were divided into three fluctuation schedules with 28 biological replicates and four blanks each. Each fluctuation schedule had a different media which changed every 80 generations. Different wells in the plates were kept blank (media without cells) to monitor for contamination. The layout of the plates and the schematic of the evolution is shown in Fig. 1A, Fig. 1B.

Daily transfers were done using a Biomek FXp robot (Beckman Coulter). Populations were diluted 1:256 daily, determining 8 generations per day on average across conditions and strains. Evolution was carried out for 39 days (a total of 312 generations). Every day the media was pipetted into 12-well reservoirs that were used to fill the plates with robotic tips FX-250-R-S (Axygen); pipette tips were reused with the same type of media. Plates were kept shaking at 1300 rpm until the transfer was performed. Six different Biomek p50 pipette tip boxes (B01090, Beckman Coulter) were used in the transfers, each corresponding to one plate. The tips were washed with water and 100% ethanol every day and left to dry overnight to reuse the following day. 96-well plates were re-used after being washed with bleach, distilled water, and autoclaved (121°C, 15 min). Every 7 days, aliquots of all the populations were frozen in 6% glycerol (final concentration) at −80°C. In case of population crashes (no growth after 24 hours from the transfer) 4 µl were transferred from the last frozen glycerol stock into 124 µl media. Throughout evolution, blank wells would occasionally be cross-contaminated, presumably from neighboring wells. When this happened, the blanks plus five random wells from the same plate were checked under the microscope for bacterial contamination. We did not detect bacterial contamination across the evolution. Due to evaporation (especially in the corners of the plate) and pipetting errors, a few populations were also lost during the experiments. A total of 274 populations were removed due to cross-contamination after genome sequence analysis (see genome analysis below).

### Fitness assays

Parental and evolved strains were revived from −80 °C glycerol stocks stored in 96-well microtiter plates. Each strain was robotically pinned (96 long-pads, Singer RoToR HDA) and arranged into a 1536 grid format on a YPD agar plate (Singer+ plate). The cultures were incubated at room temperature for four days to facilitate growth. Approximately 50,000 cells were transferred from the YPD plate to fresh Delft medium plates, using 1536 short-pads, and incubated for three days at 30°C. To minimize effects of recovery from glycerol stocks, another pinning step on YPD was made. We then pinned onto Delft plates for pre-culturing, followed by pinning onto plates containing one of 20 media conditions representative of those encountered during evolution (the 12 media used for evolution, plus an additional 8 conditions, see Table 2). The plates were then placed in the scanners for phenotyping within 30 minutes.

#### Growth Monitoring Setup

The expansion of clonal cell populations on experimental plates was monitored using the Scan-o-matic system version 2.2 (Zackrisson *et al*, 2016). Plates, without lids, were placed in high-definition desktop scanners (Epson Perfection V800 PHOTO, Epson Corporation, UK) located inside dark, humidity-controlled, and temperature-regulated (30°C) cabinets. Each scanner accommodated four plates.

#### Image Acquisition

Images were captured via transmissive scanning at 600 dpi using SANE (Scanner Access Now Easy). Plates were secured in position by an acrylic glass fixture. A transmissive grayscale calibration strip (LaserSoft IT8 Calibration Target, LaserSoft Imaging, Germany) was employed to ensure normalization and standardization of pixel intensity across different scanners and experiments. The pixel intensity of the grayscale calibration strips was benchmarked against the manufacturer’s standards, allowing for adjustment of variations in the light intensity of the transmission scans.

#### Colony Detection and Analysis

The Scan-o-matic software identified colonies using a virtual grid over each plate, aligning grid intersections with colony centers. Colonies and their surrounding areas were segmented at these intersections for local background and pixel intensity assessment. This pixel intensity data was then converted into total cell numbers using a predefined calibration function, derived from spectroscopic and flow cytometry measurements (Zackrisson *et al*, 2016).

#### Data Processing and Phenotype Extraction

Population size growth curves were extracted from these measurements. A two-step smoothing process was employed to mitigate random noise: first, local spikes were removed using a median filter; subsequently, a Gaussian filter reduced the remaining noise. These smoothed growth curves were then transformed into population doubling times and normalized biomass concentration metrics, following the methodology outlined in (Zackrisson *et al*, 2016).

### Fitness and robustness calculations

Fitness data expressed as normalized biomass and generation time were imported in R for subsequent analysis and visualization. The assessment of parental strain fitness revealed variations across replicate plates in specific measurement environments (Appendix Figure S11). Notably, the fitness distributions in formic acid 2 g/L plate 2, HMF 3 g/L plate 2, no trace metals plate 1, and NaCl plate 4 were significantly different compared to the other three plates for all two strains and were excluded from the analysis. The fitness of evolved samples was expressed as the ratio between the evolved strain and its corresponding parental strain, both revived from the same glycerol stock. For visualization purposes, the media used in the phenotypic assay were grouped according to the chemicals used (Table 2, group). Principal component analysis - PCA (prcomp function, stats library, R (STAT: Interactive Document for Working with Basic Statistical Analysis, 2019)) was used for dimensionality reduction (Fig. 2B, Fig. 3B). For better visualization of the clusters, the convex hull polygons were added to the PCA plots.

Robustness was calculated using a modified version of Eq. 3 from our previously published paper (Trivellin *et al*, 2022), where R is the robustness, σ^2^/μ is the index of dispersion and *m* is the average biomass across all environments and strains:

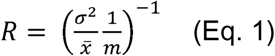

The calculation of robustness was carried out using normalized biomass and not ratio (as for the fitness). Robustness was calculated across the 20 different measurement conditions (Table 2). Each of the four replicate plates was treated as an independent data point. Robustness was outlined separately for the parental strains and the evolved populations (Fig. 3D). Differences in the distributions were assessed with a non-parametric statistical test and expressed with the p-value (wilcox.test, stat_signif function, ggsignif package R (Ahlmann-Eltze & Patil, 2022)).

### Whole-genome sequencing

Whole-genome sequencing was conducted on a total of 504 individually evolved populations, along with single colonies extracted from randomly selected wells. In the case of populations evolving in constant conditions, 48 individual colonies were sequenced for each strain, while populations evolving in fluctuating conditions had 24 single colonies sequenced for each strain. A map detailing the locations of the extracted single colonies is available in Appendix Table S5. Additionally, eight colonies from each parental strain were sequenced. Glycerol stocks from the selected wells were plated on YPD and left at 30°C for three days. Next, glycerol stocks from the final point of evolution as well as the parental strains were thawed. 10 µLfrom the stocks were inoculated in 600 µLYPD (96-well plates). Single colonies were picked from YPD plates and grown independently in 600 µLYPD as well. Cultures were left overnight at 30°C, 1200 rpm shaking. After, cultures were centrifuged at 4000 rfc for three minutes and the media was removed carefully. Pelleted cells were frozen at −20°C for DNA extraction. DNA extraction protocol was based on Bio-On-magnetic-Beads (BOMB) (Oberacker *et al*, 2019) and was performed using the Biomek FXp robot (Beckman Coulter) according to a previously setup protocol for DNA extraction (Johnson *et al*, 2021). In short, pelleted cells were resuspended in 50 µLof yeast lyses extraction buffer (5 mg/mL Zymolyase 20T (Nacalai Tesque), 1M Sorbitol, 100 mM Sodium Phosphate pH 7.4, 10 mM EDTA, 0.5% 3-(N,N-Dimethylmyristylammonio)-propanesulfonate (Sigma, T7763), 200 µg/mL RNAse A, and 20 mM DTT). The solution was left at 37°C for 45 minutes, plates were foiled to avoid evaporation. Then, 85 µLof MJ-BB buffer (modified BOMB buffer) (4M guanidinium-isothiocyanate (Goldbio G-210–500), 50 mM Tris-HCl pH 8, 20 mM EDTA) and 115 µLisopropanol (VWR# BDH1133-4LP) were added, followed by a mixing time of 5 minutes after each addition. 30 µLof magnetic beads (Zymo Research MagBinding) were added to the wells, mixed for 5 minutes and left to bind the DNA for additional 3 minutes. Magnetic beads were then separated from the rest of the solution by placing the plate on a Magnum FLX 96-well magnetic separation rack (Alpaqua) for 10 minutes while the supernatant was removed. Pellet was washed twice with 400 µLisopropanol and 300 µLof 80% ethanol. After the second wash ethanol was left to evaporate for 10 minutes. 120 µLwater were added to the pellet and mixed for 3 minutes. Finally, beads were separated from the solution and 60 µLof the supernatant were transferred into a clear 96-well plate (Bio-Rad HSP9631) for library preparation. Extracted DNA was quantified using Qubit® and a fluorescent dye (AccuGreen™ High Sensitivity dsDNA Quantitation Kit). After quantification, libraries were prepared using a Nextera (illumina) kit. Tagmentation was performed in a total volume of 2.5 µLcontaing 1 µLgDNA, 0.25 µLTDE1(Tagment DNA Enzyme, Illumina) and 1.25 µLTD buffer (Illumina). The reaction was carried out for 15 minutes at 55°C. 2.5 µLof the tagmented DNA together with 11 µLKAPA HIFi HotStart ReadyMix (Roche), 2.2 µLof each designed primers S500 and N700 (list in Appendix Table S6) and 4.1 µLddH_2_O were used in the PCR reaction (Table 3). Amplified fragments were run on an agarose gel to evaluate quality and size. Samples were pulled together based on concentration and fragment size. Samples were first pulled in 35 different tubes then into 4 different tubes. The four pools were used for a two-sided bead-based size selection (0.85-0.55) PCRClean DX Magnetic Beads (Aline). Concentrations of the pooled DNA was evaluated using Qubit® and the fragments were checked in a tape station using a D5000 ScreenTape®. Libraries were sequenced to an average depth of 20-fold using Illumina NovaSeq.

**Table 3:** PCR program.

| Temperature (°C) | time |
| --- | --- |
| 72 | 3 min |
| 98 | 5 min |
| 98 | 10 sec |
| 63 | 30 sec |
| 72 | 30 sec |
| Cycle to third step X13 times |  |
| 72 | 5 min |

### Sequencing analysis

#### Mapping and variant calling

Adapters were trimmed from the reads using Cutadapt (v 4.0) and mapped using Bwa mem (v. 0.7.17). The mapped reads were then sorted with Samtools (v 1.9) and duplicate reads were marked using MarkDuplicates from GATK (v.4.1.4.1). All strains were mapped against the S288C reference (GCF_000146045.2_R64_genomic.fna, downloaded from ncbi <u>website</u>). Variant calling was performed with Freebayes (v. 1.3.2), with −0 for stringent input base and mapping quality filters. At least 4 reads carrying the alternate allele and a minimum alternate allele fraction of 0.2 was required for the position to be evaluated (freebayes −0 -C 4 -F 0.2 -n 4). The reference genome was split and variant calling was performed on all samples together in smaller regions. All regions were concatenated using Bcftools concat (v. 1.14).

#### Contaminated samples

Some samples appeared to contain DNA from two of the parental strains. To identify these samples, we extracted 15 SNPs specific to Ethanol Red. The variant allele frequency (VAF) of these 15 SNPs were extracted for all samples, and samples with a mean VAF > 0.4 were considered Ethanol Red samples and samples with VAF < 0.02 were considered pure S288C. All other samples, as well as samples with missing data, were removed from analysis. A total of 46 S288C, and 65 Ethanol Red samples remained. In total 274 lines were removed from the analysis.

#### Variant filters

Vcftools was used to remove variants with a quality < 20, mean depth < 10 or > 100, missingness >5% as well as variants with more than two alleles. All genotypes with < 5 reads coverage were excluded as well as variants with an allele count ≥ 1 in at least 10 samples. Variants were annotated using Variant Effect Predictor (110.1).

#### Counting mutations in each evolution regime

The variant allele frequency was extracted for all remaining variants and variants with a variant allele count ≥ 5 and VAF > 0.4 (Ethanol Red) or VAF >0.8 (S288C) in at most 4 samples were considered potential (fixed) novel mutations (Table 4).

**Table 4:** count of unique variants in each strain.

| Strain | n samples | Novel variants |
| --- | --- | --- |
| S288C | 46 | 83 |
| Ethanol Red | 65 | 65 |

### Statistical enrichment analysis

Single colonies variants were used as input for the enrichment analysis that led to identifications of gene ontology terms, biological pathways, regulatory motifs, transcription factors, cellular compartment, and pathways. The analysis was performed with the gprofiler2 package in R using the gost, gostplot and publish_gosttable functions with g:SCS for multiple testing correction for p-value (Kolberg *et al*, 2023).

### Analysis of multi-hit genes and parallelism

To investigate parallelism in the set of unique variants, only the non-synonymous mutations were considered (synonymous, intergenic, splice region, non-coding exon and in frame deletions variants were filtered out). A complete list of genes from the *Saccharomyces cerevisiae* S288C strain was downloaded from the NIH national library of medicine containing 6022 genes in total (https://www.ncbi.nlm.nih.gov/gene - txid 559292). Population hits were calculated as the number of hits in each gene across populations. Multiplicity was defined as the population hits divided by the length of each gene (base pairs) multiplied by the mean gene length across the whole sets of genes in the S288C genome. The null model was built by assigning the hits from our data to random genes from the S288C genome weighted by the gene length. 1000 simulations were drawn to generate null distributions both for the multiplicity and the population hits. To assess if the distribution of the population hits and multiplicity were significantly different from the null distributions, a nonparametric statistical test (Mann–Whitney U test) was used and a p-value was calculated (Appendix Figure S9).

## Declarations

### Author contributions

CT conceived the idea, planned and carried out the evolution experiments and the library preparation for sequencing. MG helped with DNA extractions, library preparation and genome sequencing. CT and KP performed the phenotypic assays. KP performed the pre-processing of the phenotypic data. CT performed the data analysis of fitness and robustness. DE performed genome sequencing analysis. CT performed data analysis from the variants data. CT, together with MD and LO, interpreted the results. CT wrote the manuscript, DE wrote the genome sequence analysis materials and methods, KP wrote the materials and methods for the phenotypic assays. CT edited the manuscript together with all the other authors.

## Acknowledgements

Financial support by Novo Nordisk Foundation Grant Distinguished Investigator 2019 -Research within biotechnology-based synthesis & production (#0055044) is gratefully acknowledged. We would like to express our gratitude to Société Industrielle Lesaffre, Division Leaf, for providing the Ethanol Red strain. The Bauer Core Facility at Harvard University and Bioinformatics Long-Term Support at SciLifeLab (Science for Life Laboratory, Sweden) are kindly acknowledged for their assistance with sequencing and genomic analysis. The computations and data handling were enabled by resources provided by the National Academic Infrastructure for Supercomputing in Sweden (NAISS), partially funded by the Swedish Research Council through grant agreement no. 2022-06725. Acknowledgment is given to the Chalmers Energy Area of Advance and the Barbro Osher Pro Suecia Foundation for their support in internationalization efforts and funding for research visits.

## Data availability

The data for this study have been deposited in the European Nucleotide Archive (ENA) at EMBL-EBI under accession number PRJEB104644.

Fitness data and scripts to analyze fitness, robustness and variants can be find here: github.com/cectri/evolution_robustness_yeast.

## Appendix

Appendix Figure S1-S11

Appendix Table S1-S6

## References

Abdul-Rahman F, Tranchina D & Gresham D (2021) Fluctuating Environments Maintain Genetic Diversity through Neutral Fitness Effects and Balancing Selection. Mol Biol Evol 38: 4362–4375

Ahlmann-Eltze C & Patil I (2022) Significance Brackets for ‘ggplot2’ [R package ggsignif version 0.6.4]. CRAN: Contributed Packages

Bouyx C, Schiavone M & François JM (2021) FLO11, a Developmental Gene Conferring Impressive Adaptive Plasticity to the Yeast Saccharomyces cerevisiae. Pathogens 10

Bruinenberg PM, Van Dijken JP & Scheffers WA (1983) An enzymic analysis of NADPH production and consumption in Candida utilis. J Gen Microbiol 129: 965–971

Chandler CH, Chari S, Tack D & Dworkin I (2014) Causes and consequences of genetic background effects illuminated by integrative genomic analysis. Genetics 196: 1321–1336

Chen V, Johnson MS, Hérissant L, Humphrey PT, Yuan DC, Li Y, Agarwala A, Hoelscher SB, Petrov DA, Desai MM, et al (2023) Evolution of haploid and diploid populations reveals common, strong, and variable pleiotropic effects in non-home environments. Elife 12: 92899

Cooper TF & Lenski RE (2010) Experimental evolution with E. coli in diverse resource environments. I. Fluctuating environments promote divergence of replicate populations. BMC Evol Biol 10: 1–10

Cvijović I, Good BH, Jerison ER & Desai MM (2015) Fate of a mutation in a fluctuating environment. Proc Natl Acad Sci U S A 112: E5021–E5028

Dhar R, Sägesser R, Weikert C & Wagner A (2013) Yeast adapts to a changing stressful environment by evolving cross-protection and anticipatory gene regulation. Mol Biol Evol 30: 573–588

Dolpatcha S, Phong HX, Thanonkeo S, Klanrit P, Yamada M & Thanonkeo P (2023) Adaptive laboratory evolution under acetic acid stress enhances the multistress tolerance and ethanol production efficiency of Pichia kudriavzevii from lignocellulosic biomass. Scientific Reports 2023 13:1 13: 1–15

Dragosits M & Mattanovich D (2013) Adaptive laboratory evolution -principles and applications for biotechnology. Microb Cell Fact 12: 1–17

Félix M-AA & Barkoulas M (2015) Pervasive robustness in biological systems. Nature Reviews Genetics 2015 16:8 16: 483–496

Fernández-Vázquez J, Vargas-Pérez I, Sansó M, Buhne K, Carmona M, Paulo E, Hermand D, Rodríguez-Gabriel M, Ayté J, Leidel S, et al (2013) Modification of tRNALys UUU by Elongator Is Essential for Efficient Translation of Stress mRNAs. PLoS Genet 9: e1003647

Goldstein I & Ehrenreich IM (2021) The complex role of genetic background in shaping the effects of spontaneous and induced mutations. Yeast 38: 187

Gong A, Liu W, Lin Y, Huang L & Xie Z (2023) Adaptive Laboratory Evolution Reveals the Selenium Efflux Process To Improve Selenium Tolerance Mediated by the Membrane Sulfite Pump in Saccharomyces cerevisiae. Microbiol Spectr 11

Gonzalez A & Bell G (2013) Evolutionary rescue and adaptation to abrupt environmental change depends upon the history of stress. Philosophical Transactions of the Royal Society B: Biological Sciences 368

Good BH, McDonald MJ, Barrick JE, Lenski RE & Desai MM (2017) The Dynamics of Molecular Evolution Over 60,000 Generations. Nature 551: 45

Gronchi N, De Bernardini N, Cripwell RA, Treu L, Campanaro S, Basaglia M, Foulquié-Moreno MR, Thevelein JM, Van Zyl WH, Favaro L, et al (2022) Natural Saccharomyces cerevisiae Strain Reveals Peculiar Genomic Traits for Starch-to-Bioethanol Production: the Design of an Amylolytic Consolidated Bioprocessing Yeast. Front Microbiol 12: 768562

Haaland TR, Wright J & Ratikainen II (2020) Generalists versus specialists in fluctuating environments: a bet-hedging perspective. Oikos 129: 879–890

Ikner A & Shiozaki K (2005) Yeast signaling pathways in the oxidative stress response. Mutation Research/Fundamental and Molecular Mechanisms of Mutagenesis 569: 13–27

Johnson MS, Gopalakrishnan S, Goyal J, Dillingham ME, Bakerlee CW, Humphrey PT, Jagdish T, Jerison ER, Kosheleva K, Lawrence KR, et al (2021) Phenotypic and molecular evolution across 10,000 generations in laboratory budding yeast populations. Elife 10: 1–28

Kang K, Bergdahl B, MacHado D, Dato L, Han TL, Li J, Villas-Boas S, Herrgård MJ, Förster J & Panagiotou G (2019) Linking genetic, metabolic, and phenotypic diversity among Saccharomyces cerevisiae strains using multi-omics associations. Gigascience 8: 1–14

Ketola T, Mikonranta L, Zhang J, Saarinen K, Örmälä AM, Friman VP, Mappes J & Laakso J (2013) FLUCTUATING TEMPERATURE LEADS TO EVOLUTION OF THERMAL GENERALISM AND PREADAPTATION TO NOVEL ENVIRONMENTS. Evolution (N Y) 67: 2936–2944

Kim J & Rose MD (2015) Stable Pseudohyphal Growth in Budding Yeast Induced by Synergism between Septin Defects and Altered MAP-kinase Signaling. PLoS Genet 11: e1005684

Kitano H (2004) Biological robustness. Nat Rev Genet: 826–837 doi:10.1038/nrg1471 [PREPRINT]

Kolberg L, Raudvere U, Kuzmin I, Adler P, Vilo J & Peterson H (2023) g:Profiler—interoperable web service for functional enrichment analysis and gene identifier mapping (2023 update). Nucleic Acids Res 51: W207–W212

Kvitek DJ & Sherlock G (2013) Whole Genome, Whole Population Sequencing Reveals That Loss of Signaling Networks Is the Major Adaptive Strategy in a Constant Environment. PLoS Genet 9: e1003972

Levy SF & Siegal ML (2008) Network Hubs Buffer Environmental Variation in Saccharomyces cerevisiae. PLoS Biol 6: e264

Mavrommati M, Daskalaki A, Papanikolaou S & Aggelis G (2022) Adaptive laboratory evolution principles and applications in industrial biotechnology. Biotechnol Adv 54

Menegon YA, Gross J & Jacobus AP (2022) How adaptive laboratory evolution can boost yeast tolerance to lignocellulosic hydrolyses. Current Genetics 2022 68:3 68: 319–342

Mohler K, Mann R, Kyle A, Reynolds N & Ibba M (2018) Aminoacyl-tRNA quality control is required for efficient activation of the TOR pathway regulator Gln3p. RNA Biol 15: 594–603

Ndukwe JK, Aliyu GO, Onwosi CO, Chukwu KO & Ezugworie FN (2020) Mechanisms of weak acid-induced stress tolerance in yeasts: Prospects for improved bioethanol production from lignocellulosic biomass. Process Biochemistry 90: 118–130

Oberacker P, Stepper P, Bond DM, Höhn S, Focken J, Meyer V, Schelle L, Sugrue VJ, Jeunen GJ, Moser T, et al (2019) Bio-On-Magnetic-Beads (BOMB): Open platform for high-throughput nucleic acid extraction and manipulation. PLoS Biol 17: e3000107

Olsson L, Rugbjerg P, Torello Pianale L & Trivellin C (2022) Robustness: linking strain design to viable bioprocesses. Trends Biotechnol

Pereira R, Wei Y, Mohamed E, Radi M, Malina C, Herrgård MJ, Feist AM, Nielsen J & Chen Y (2019) Adaptive laboratory evolution of tolerance to dicarboxylic acids in Saccharomyces cerevisiae. Metab Eng 56: 130–141

Pérez-Gallardo R V., Briones LS, Díaz-Pérez AL, Gutiérrez S, Rodríguez-Zavala JS & Campos-García J (2013) Reactive oxygen species production induced by ethanol in Saccharomyces cerevisiae increases because of a dysfunctional mitochondrial iron–sulfur cluster assembly system. FEMS Yeast Res 13: 804–819

Persson K, Stenberg S, Tamás MJ & Warringer J (2022) Adaptation of the yeast gene knockout collection is near-perfectly predicted by fitness and diminishing return epistasis. G3 Genes|Genomes|Genetics 12

Sánchez-Adriá IE, Sanmartín G, Prieto JA, Estruch F, Fortis E & Randez-Gil F (2023) Adaptive laboratory evolution for acetic acid-tolerance matches sourdough challenges with yeast phenotypes. Microbiol Res 277: 127487

Sandberg TE, Salazar MJ, Weng LL, Palsson BO & Feist AM (2019) The emergence of adaptive laboratory evolution as an efficient tool for biological discovery and industrial biotechnology. Metab Eng 56: 1

Sangster TA, Lindquist S & Queitsch C (2004) Under cover: Causes, effects and implications of Hsp90-mediated genetic capacitance. BioEssays 26: 348–362

Shivaswamy S & Iyer VR (2008) Stress-Dependent Dynamics of Global Chromatin Remodeling in Yeast: Dual Role for SWI/SNF in the Heat Shock Stress Response. Mol Cell Biol 28: 2221

STAT: Interactive Document for Working with Basic Statistical Analysis (2019) CRAN: Contributed Packages

Szklarczyk D, Kirsch R, Koutrouli M, Nastou K, Mehryary F, Hachilif R, Gable AL, Fang T, Doncheva NT, Pyysalo S, et al (2023) The STRING database in 2023: protein–protein association networks and functional enrichment analyses for any sequenced genome of interest. Nucleic Acids Res 51: D638–D646

Thompson DA, Desai MM & Murray AW (2006) Ploidy Controls the Success of Mutators and Nature of Mutations during Budding Yeast Evolution. Current Biology 16: 1581–1590

Torello Pianale L, Caputo F & Olsson L (2023) Four ways of implementing robustness quantification in strain characterisation. Biotechnology for Biofuels and Bioproducts 16: 1–20

Trivellin C, Olsson L & Rugbjerg P (2022) Quantification of Microbial Robustness in Yeast. ACS Synth Biol 11: 1686–1691

Trivellin C, Rugbjerg P & Olsson L (2024a) Performance and robustness analysis reveals phenotypic trade-offs in yeast. Life Sci Alliance 7: e202302215

Trivellin C, Torello Pianale L & Olsson L (2024b) Robustness quantification of a mutant library screen revealed key genetic markers in yeast. Microb Cell Fact 23

Wallace-Salinas V, Brink DP, Ahrén D & Gorwa-Grauslund MF (2015) Cell periphery-related proteins as major genomic targets behind the adaptive evolution of an industrial Saccharomyces cerevisiae strain to combined heat and hydrolysate stress. BMC Genomics 16: 1–16

Walsh P, Bursać D, Law YC, Cyr D & Lithgow T (2004) The J-protein family: Modulating protein assembly, disassembly and translocation. EMBO Rep 5: 567–571

Wenger JW, Piotrowski J, Nagarajan S, Chiotti K, Sherlock G & Rosenzweig F (2011) Hunger Artists: Yeast Adapted to Carbon Limitation Show Trade-Offs under Carbon Sufficiency. PLoS Genet 7: e1002202

Wiser MJ, Ribeck N & Lenski RE (2013) Long-term dynamics of adaptation in asexual populations. Science 342: 1364–1367

Zackrisson M, Hallin J, Ottosson LG, Dahl P, Fernandez-Parada E, Ländström E, Fernandez-Ricaud L, Kaferle P, Skyman A, Stenberg S, et al (2016) Scan-o-matic: High-resolution microbial phenomics at a massive scale. G3: Genes, Genomes, Genetics 6: 3003–3014

