## Appendix for "Impact of fluctuating environments on the fitness and robustness of evolving laboratory and industrial *Saccharomyces cerevisiae* strains"

**Table of contents:**

### Appendix Figures S1–S11

### Appendix Tables S1-S6

**Appendix figures**

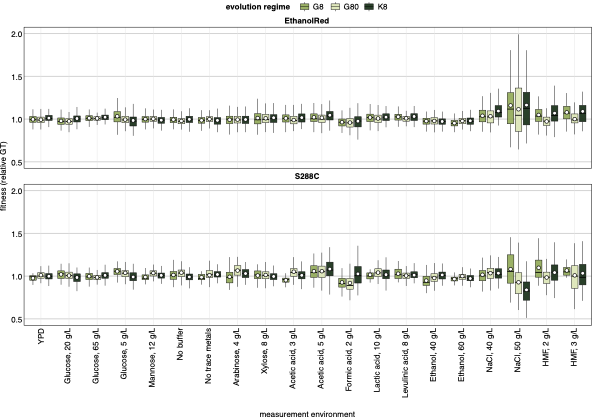

**Appendix Figure S1 - Comparison of fitness (relative generation time) under different evolution regimes.**

Data information: The relative generation time (GT of each evolved strain divided by the GT of its parental strain) is shown on the y-axis. Each boxplot represents the distribution of relative GT for populations evolved under a given regime (colors). The black horizontal line indicates the median, and the white dot indicates the mean. Distributions are shown across measurement environments (x-axis), evolution regimes (color), and strains (facets).

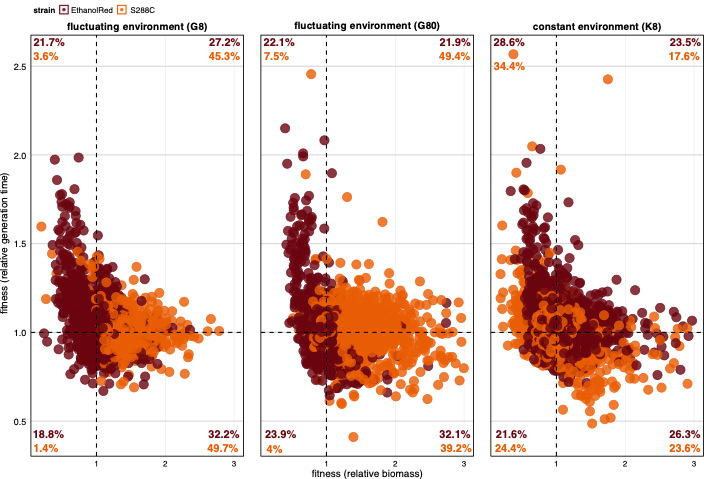

**Appendix Figure S2 - Comparison of generation time changes versus biomass changes**.

Data information: The relative generation time (GT of the evolved strains / GT of the parental strain, y-axis) is plotted versus the relative biomass (biomass of the evolved strains / biomass of the parental strain, x-axis). Each dot corresponds to a single population and is colored by strain. Comparison is made for each evolution regime (facets). The two dotted lines, vertical and horizontal, represent no change in fitness compared to the parentals. The percentages in each quadrant represent the percentage of populations (of each strain showed by the color) that are in that quadrant compared to the total number of each strain in each facet.

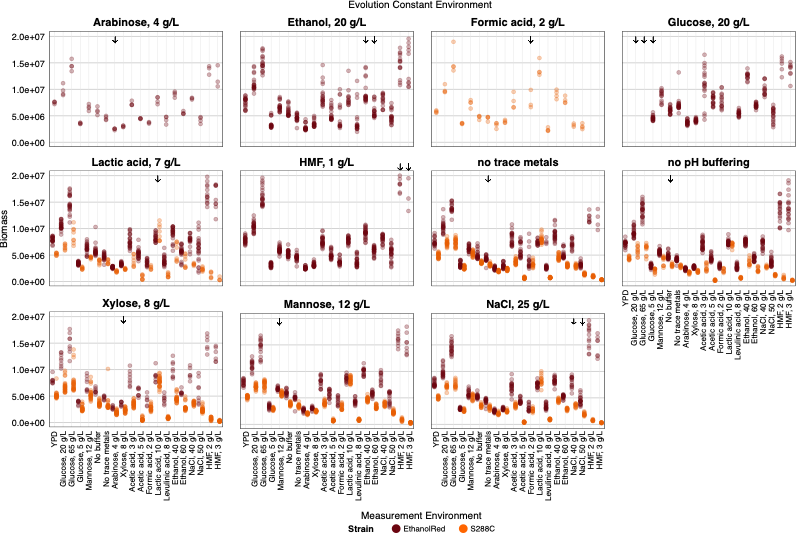

**Appendix Figure S3 - Biomass of evolved K8 populations.**

Data information: The biomass of each population evolving in constant environments is shown on the y-axis for each evolution environment (facets). The x-axis shows the measurement environments. Strains are shown in different colors. Arrows represent the fitness in the ´home´ environment.

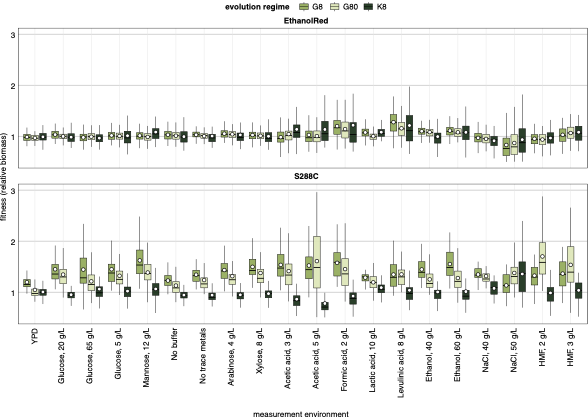

**Appendix Figure S4 - Comparison of fitness (relative biomass) under different evolution regimes**.

Data information: The relative biomass (biomass of each evolved strain divided by the biomass of the corresponding parental strain) is shown on the y-axis. Each boxplot represents the distribution of relative biomass for populations evolved under a given regime (colors). The black horizontal line indicates the median, and the white dot corresponds the mean. Distributions are shown across measurement environments (x-axis), evolution regimes (color), and strains (facets).

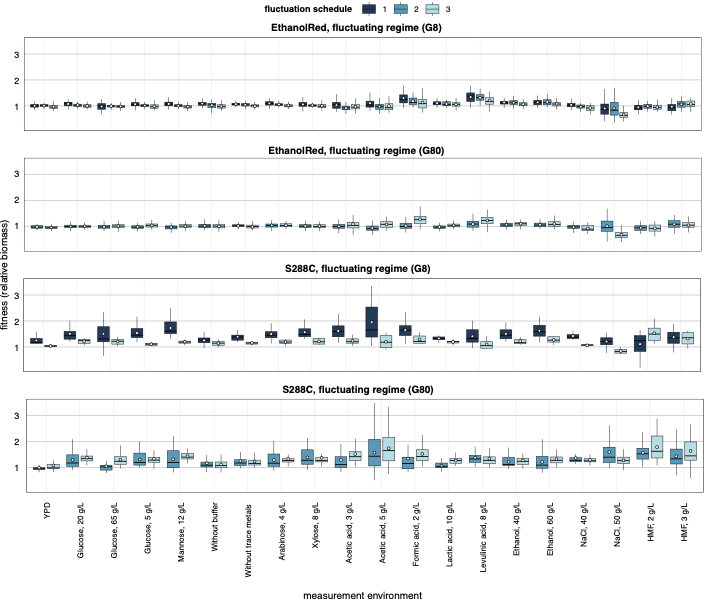

**Appendix Figure S5 - Comparison of fitness (relative biomass) under different fluctuation schedules.**

Data information: The relative fitness (biomass of the evolved strains / biomass of the matching parental strain) is plotted on the y-axis. Each boxplot corresponds to the distribution of populations evolved in fluctuating regimes only and is plotted for each measurement environment (x-axis), evolution regime and strain (facet), and fluctuation schedule (color). Missing data are due to contamination issues (see Materials and Methods).

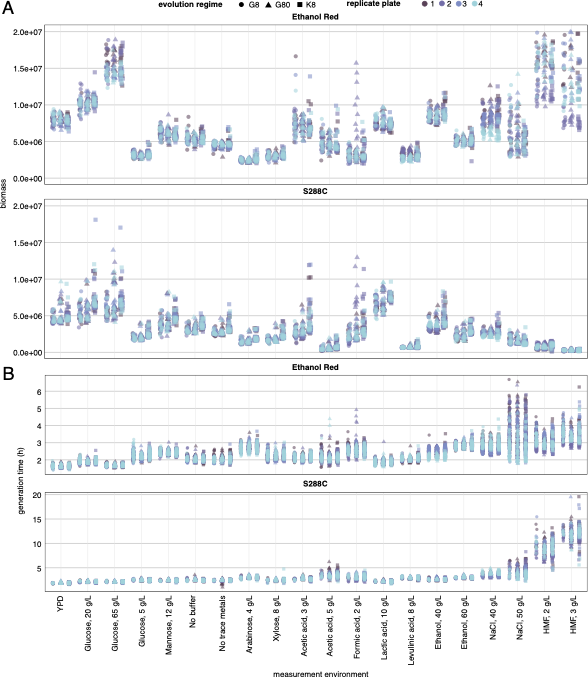

**Appendix Figure S6 - Fitness of parental strains.**

**A** The biomass of parental strains is shown on the y-axis for each evolution regime (shape) and each replicate plate from the scan-o-matic assay (color). The x-axis shows the measurement environments. Each facet represents one strain.

**B** as in **A** but the generation time (h) of the parental strains is shown on the y-axis instead of the biomass.

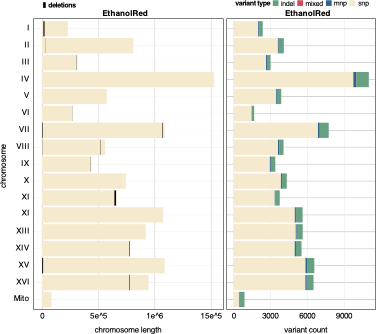

**Appendix Figure S7 - Genomic variants analysis of Ethanol Red compared to the S288C reference genome.**

Data information: In the left plot, chromosomes are plotted on the y-axis. Deletions in the parental Ethanol Red strain are represented by the black vertical line in each chromosome. The list of parental deletion can be found in Appendix Table S1. In the right plot, variant count (x-axis) divided by type (colors), is plotted for each chromosome (y-axis).

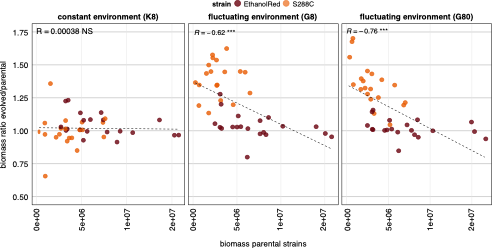

**Appendix Figure S8 - Correlation between parental fitness and fitness gain at the final evolution time point.**

Data information: The biomass of the parental strains is shown on the x-axis and the relative biomass (evolved/parental) is plotted on the y-axis. Each dot corresponds to the average measurement across populations and replicates for each measurement environment (n=20). The dots are colored by strain. The correlations are divided by evolution regime (facets). The dotted line shows the trend of the spearman correlation between x and y and the coefficient is shown on the top left corner together with stars indicating the significance, ´***´means p-value < 0.001 and ´NS´ means not significant p-value >0.05.

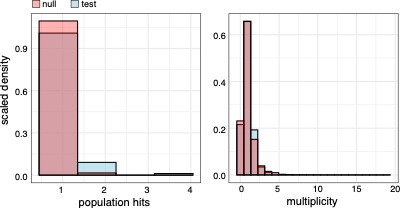

**Appendix Figure S9 - Parallelism.**

Data information: Comparison of test and null distribution of population hits (left) and multiplicity (right) (see Materials and Methods for specifics of the simulations).

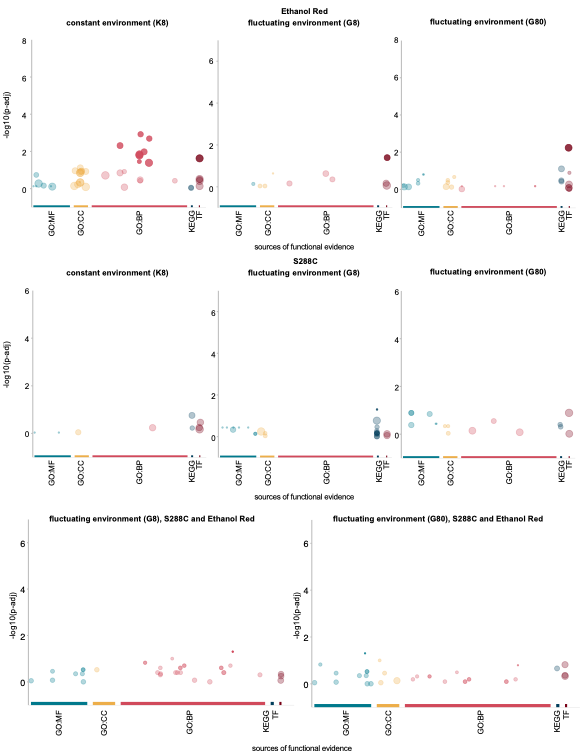

**Appendix Figure S10 - Gene Ontology (GO) analysis.**

Data information: The analysis of the GO terms was divided by evolution regime and strains (name of each plot). GO terms are shown by the colored dots. The less transparent dots represent the significantly enriched terms (padj < 0.05). The y-axis shows the -log10(padj). The x-axis shows different categories of GO-terms (MF: molecular function; CC: cellular component; BP: biological process; KEGG: Kegg pathways; TF: transcription factors). Corresponding GO:ID and terms are explained in Appendix Table S2. Plots were generated using g:Profiler (see Materials and Methods).

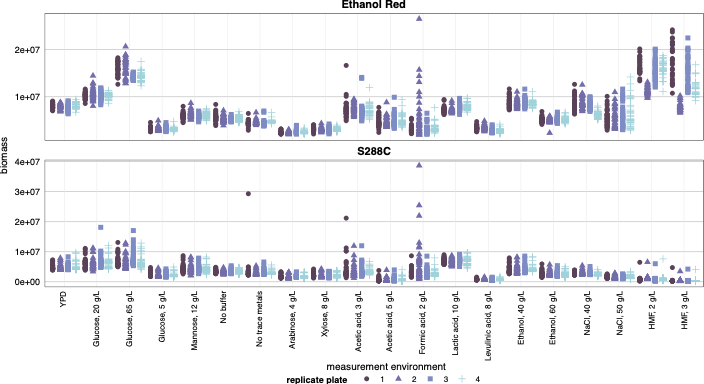

**Appendix Figure S11 - Fitness assay replicate plates assessment**.

Data information: The biomass of parental strains is shown on the y-axis for each replicate plate from the scan-o-matic assay (color and shape) (see Materials and Methods). The x-axis shows the measurement environments. Each facet represents one strain.

**Appendix Tables**

**Appendix Table S1.** List of deletions identified in parental strain Ethanol Red compared to the S288C reference genome. Deletion position within the chromosomes and affected genes are reported.

| **start** | **end** | **strain** | **length** | **chromosome** | **systematic name** | **gene name** |
| --- | --- | --- | --- | --- | --- | --- |
| 13650 | 23675 | EthanolRed | 10025 | I | YAL064C-A | TDA8 |
| 13650 | 23675 | EthanolRed | 10025 | I | YAL064W | YAL064W |
| 13650 | 23675 | EthanolRed | 10025 | I | YAL063C-A | YAL063C-A |
| 32625 | 34025 | EthanolRed | 1400 | II | YBL100W-B | YBL100W-B |
| 303815 | 305835 | EthanolRed | 2020 | III | YCR102C | YCR102C |
| 268747 | 270158 | EthanolRed | 1411 | VI | YFR057W | YFR057W |
| 1138 | 6301 | EthanolRed | 5163 | VII | YGL263W | COS12 |
| 1138 | 6301 | EthanolRed | 5163 | VII | YGL262W | YGL262W |
| 1138 | 6301 | EthanolRed | 5163 | VII | YGL261C | PAU11 |
| 1071675 | 1076148 | EthanolRed | 4473 | VII | YGR288W | MAL13 |
| 1071675 | 1076148 | EthanolRed | 4473 | VII | YGR289C | MAL11 |
| 1411 | 3320 | EthanolRed | 1909 | VIII | YHL050C | YHL050C |
| 522456 | 525443 | EthanolRed | 2987 | VIII | YHR211W | FLO5 |
| 433795 | 435392 | EthanolRed | 1597 | IX | YIR041W | PAU15 |
| 433795 | 435392 | EthanolRed | 1597 | IX | YIR042C | YIR042C |
| 645199 | 659136 | EthanolRed | 13937 | XI | YKR102W | FLO10 |
| 645199 | 659136 | EthanolRed | 13937 | XI | YKR104W | YKR104W |
| 645199 | 659136 | EthanolRed | 13937 | XI | YKR105C | VBA5 |
| 645199 | 659136 | EthanolRed | 13937 | XI | YKR103W | NFT1 |
| 776858 | 781290 | EthanolRed | 4432 | XIV | YNR074C | AIF1 |
| 776858 | 781290 | EthanolRed | 4432 | XIV | YNR075W | COS10 |
| 2558 | 11104 | EthanolRed | 8546 | XV | YOL164W-A | YOL164W-A |
| 2558 | 11104 | EthanolRed | 8546 | XV | YOL164W | BDS1 |
| 2558 | 11104 | EthanolRed | 8546 | XV | YOL163W | YOL163W |
| 2558 | 11104 | EthanolRed | 8546 | XV | YOL162W | YOL162W |
| 777214 | 781621 | EthanolRed | 4407 | XVI | YPR121W | THI22 |

**Appendix Table S2.** Significant Gene Ontology (GO) terms are summarized in each row. GO term source, ID and name are specified in the different columns and the p_adj_ is shown in the last column. The test was performed using “g:profiler” (see Materials and Methods).

| **Evolution Regime** | **strain** | **source** | **TERM ID** | **TERM NAME** | **p_value** |
| --- | --- | --- | --- | --- | --- |
| G8 | S288C and Ethanol Red | GO:BP | GO:1900187 | regulation of cell adhesion involved in single−species biofilm formation | 0.05 |
| G8 | S288C and Ethanol Red | GO:BP | GO:1900189 | positive regulation of cell adhesion involved in single−species biofilm formation | 0.05 |
| G80 | S288C and Ethanol Red | GO:MF | GO:0106261 | tRNA uridine(34) acetyltransferase activity | 0.05 |
| G8 | Ethanol Red | TF | TF:M01810_1 | Factor: Tec1p; motif: RMATTCYY; match class: 1 | 0.044 |
| G80 | Ethanol Red | TF | TF:M01901 | Factor: Abf1p; motif: YCGTNNNNNRTGAYNN | 0.0061 |
| K8 | Ethanol Red | GO:BP | GO:0051223 | regulation of protein transport | 0.0026 |
| K8 | Ethanol Red | GO:BP | GO:0070201 | regulation of establishment of protein localization | 0.0041 |
| K8 | Ethanol Red | GO:BP | GO:0032880 | regulation of protein localization | 0.0075 |
| K8 | Ethanol Red | GO:BP | GO:0050794 | regulation of cellular process | 0.016 |
| K8 | Ethanol Red | GO:BP | GO:0060341 | regulation of cellular localization | 0.018 |
| K8 | Ethanol Red | GO:BP | GO:0050789 | regulation of biological process | 0.019 |
| K8 | Ethanol Red | GO:BP | GO:0050708 | regulation of protein secretion | 0.039 |
| K8 | Ethanol Red | GO:BP | GO:0065007 | biological regulation | 0.046 |
| K8 | Ethanol Red | TF | TF:M01520 | Factor: Rsc30p; motif: NNNNNCGCGCGCGSGNNNSNN | 0.026 |
| G8 | S288C | KEGG | KEGG:00511 | Other glycan degradation | 0.05 |

**Appendix Table S3.** Composition of the trace metals solution.

| **Chemical** | **Amount (g/L)** |
| --- | --- |
| EDTA | 0.015 |
| ZnSO_4_·7H_2_O | 0.0045 |
| MnCl_2_·4H_2_O | 0.0008 |
| CoCl_2_·6H_2_O | 0.0003 |
| CuSO_4_·5H_2_O | 0.0003 |
| Na_2_MoO_4_·2H_2_O | 0.0004 |
| CaCl_2_·2H_2_O | 0.0045 |
| FeSO_4_·7H_2_O | 0.003 |
| H_3_BO_3_ | 0.001 |
| KI | 0.0001 |

**Appendix Table S4.** Composition of the vitamin solution.

| **Vitamin** | **Amount (g/L)** |
| --- | --- |
| d-Biotin | 0.00005 |
| Calcium D(+) pantothenate | 0.001 |
| Nicotinic acid | 0.001 |
| Myo-inositol | 0.025 |
| Thiamine HCl | 0.001 |
| Pyridoxine HCl | 0.001 |
| Para-aminobenzoic acid | 0.0002 |

**Appendix Table S5**. Scheme of the single colonies isolated from the evolution plates and grown in separate 96-well plates (see Materials and Methods).

(separate file)

**Appendix Table S6.** Forward and Reverse primers used for fragment amplification during library preparation (see Materials and Methods).

(separate file)
