## Appendix Table S5 for "Impact of fluctuating environments on the fitness and robustness of evolving laboratory and industrial *Saccharomyces cerevisiae* strains"

Layout of single colonies picked from evolved populations grown onYPD agar and grown in 96-well plates with YPD

PLATE1_COLONIES

| | 1 | 2 | 3 | 4 | 5 | 6 | 7 | 8 | 9 | 10 | 11 | 12 |
|---|---|---|---|---|---|---|---|---|---|---|---|---|
| A | SCB1_1 | SCA3_1 | SCB5_1 | SCC7_1 | SCB9_1 | SCG11_1 | | | | | | |
| B | SCB1_2 | SCA3_2 | SCB5_2 | SCC7_2 | SCB9_2 | SCG11_2 | | | | | | |
| C | SCE1_1 | SCG3_1 | SCG5_1 | SCE7_1 | SCF9_1 | SCC11_1 | | | | | | |
| D | SCE1_2 | SCG3_2 | SCG5_2 | SCE7_2 | SCF9_2 | SCC11_2 | | | | | | |
| E | SCD2_1 | SCC4_1 | SCA6_1 | SCD8_1 | SCA10_1 | SCB12_1 | | | | | | |
| F | SCD2_2 | SCC4_2 | SCA6_2 | SCD8_2 | SCA10_2 | SCB12_2 | | | | | | |
| G | SCF2_1 | SCE4_1 | SCF6_1 | SCG8_1 | SCD10_1 | SCF12_1 | | | | | | |
| H | SCF2_2 | SCE4_2 | SCF6_2 | SCG8_2 | SCD10_2 | SCF12_2 | | | | | | |

PLATE2_COLONIES

| | 1 | 2 | 3 | 4 | 5 | 6 | 7 | 8 | 9 | 10 | 11 | 12 |
|---|---|---|---|---|---|---|---|---|---|---|---|---|
| A | ECB1_1 | ECA3_1 | ECB5_1 | ECC7_1 | ECB9_1 | ECC11_1 | S1E1_1 | S1F5_1 | S1D9_1 | | | |
| B | ECB1_2 | ECA3_2 | ECB5_2 | ECC7_2 | ECB9_2 | ECC11_2 | S1E1_2 | S1F5_2 | S1D9_2 | | | |
| C | ECE1_1 | ECG3_1 | ECG5_1 | ECE7_1 | ECH9_1 | ECG11_1 | S1B2_1 | S1C6_1 | S1G10_1 | | | |
| D | ECE1_2 | ECG3_2 | ECG5_2 | ECE7_2 | ECH9_2 | ECG11_2 | S1B2_2 | S1C6_2 | S1G10_2 | | | |
| E | ECH2_1 | ECC4_1 | ECA6_1 | ECD8_1 | ECA10_1 | ECB12_1 | S1G3_1 | S1H7_1 | S1A11_1 | | | |
| F | ECH2_2 | ECC4_2 | ECA6_2 | ECD8_2 | ECA10_2 | ECB12_2 | S1G3_2 | S1H7_2 | S1A11_2 | | | |
| G | ECD2_1 | ECE4_1 | ECH6_1 | ECG8_1 | ECD10_1 | ECH12_1 | S1A4_1 | S1B8_1 | S1C12_1 | | | |
| H | ECD2_2 | ECE4_2 | ECH6_2 | ECG8_2 | ECD10_2 | ECH12_2 | S1A4_2 | S1B8_2 | S1C12_2 | | | |

PLATE3_COLONIES

| | 1 | 2 | 3 | 4 | 5 | 6 | 7 | 8 | 9 | 10 | 11 | 12 |
|---|---|---|---|---|---|---|---|---|---|---|---|---|
| A | E1E1_1 | E1F5_1 | E1D9_1 | S2E1_1 | S2F5_1 | S2D9_1 | | | | E2E2_1 | E2F5_1 | E2D9_1 |
| B | E1E1_2 | E1F5_2 | E1D9_2 | S2E1_2 | S2F5_2 | S2D9_2 | | | | E2E2_2 | E2F5_2 | E2D9_2 |
| C | E1B2_1 | E1C6_1 | E1G10_1 | S2G3_1 | S2C6_1 | S2G10_1 | | | | E2B2_1 | E2C6_1 | E2G10_1 |
| D | E1B2_2 | E1C6_2 | E1G10_2 | S2G3_2 | S2C6_2 | S2G10_2 | | | | E2B2_2 | E2C6_2 | E2G10_2 |
| E | E1G3_1 | E1H7_1 | E1A11_1 | S2B2_1 | S2H7_1 | S2A11_1 | | | | E2G3_1 | E2H7_1 | E2A11_1 |
| F | E1G3_2 | E1H7_2 | E1A11_2 | S2B2_2 | S2H7_2 | S2A11_2 | | | | E2G3_2 | E2H7_2 | E2A11_2 |
| G | E1A4_1 | E1B8_1 | E1C12_1 | S2A4_1 | S2B8_1 | S2C12_1 | | | | E2A4_1 | E2B8_1 | E2C12_1 |
| H | E1A4_2 | E1B8_2 | E1C12_2 | S2A4_2 | S2B8_2 | S2C12_2 | | | | E2A4_2 | E2B8_2 | E2C12_2 |

PLATE_4 colonies taken from parental strains grown on YPD agar

| | 1 | 2 | 3 |
|---|---|---|---|
| A | S1 | | E1 |
| B | S2 | | E2 |
| C | S3 | | E3 |
| D | S4 | | E4 |
| E | S5 | | E5 |
| F | S6 | | E6 |
| G | S7 | | E7 |
| H | S8 | | E8 |

| | |
|---|---|
| S= | S288C |
| E= | ETRED |

| | |
|---|---|
| C= | controls |
| 1= | G8 |
| 2= | G80 |

| | |
|---|---|
| _1= | colony 1 |
| _2= | colony2 |

B1;E1....etc wells where the colonies were isolated from
