## Appendix Table S6 for "Impact of fluctuating environments on the fitness and robustness of evolving laboratory and industrial *Saccharomyces cerevisiae* strains"

| Sample Name | i7 Index *First index read* | i5 Index *Second index read* |
|---|---|---|
| 1_S288C_controls_A1 | TAAGGCGA | GCGATCTA |
| 1_S288C_controls_A2 | TAAGGCGA | ATAGAGAG |
| 1_S288C_controls_A3 | TAAGGCGA | AGAGGATA |
| 1_S288C_controls_A4 | TAAGGCGA | TCTACTCT |
| 1_S288C_controls_A5 | TAAGGCGA | CTCCTTAC |
| 1_S288C_controls_A6 | TAAGGCGA | TATGCAGT |
| 1_S288C_controls_A7 | TAAGGCGA | TACTCCTT |
| 1_S288C_controls_A8 | TAAGGCGA | AGGCTTAG |
| 1_S288C_controls_A9 | TAAGGCGA | GAGTAGCC |
| 1_S288C_controls_A10 | TAAGGCGA | GTCTGAGG |
| 1_S288C_controls_A11 | TAAGGCGA | CGTAAGGA |
| 1_S288C_controls_A12 | TAAGGCGA | CCACGCGT |
| 1_S288C_controls_B1 | CGTACTAG | GCGATCTA |
| 1_S288C_controls_B2 | CGTACTAG | ATAGAGAG |
| 1_S288C_controls_B3 | CGTACTAG | AGAGGATA |
| 1_S288C_controls_B4 | CGTACTAG | TCTACTCT |
| 1_S288C_controls_B5 | CGTACTAG | CTCCTTAC |
| 1_S288C_controls_B6 | CGTACTAG | TATGCAGT |
| 1_S288C_controls_B7 | CGTACTAG | TACTCCTT |
| 1_S288C_controls_B8 | CGTACTAG | AGGCTTAG |
| 1_S288C_controls_B9 | CGTACTAG | GAGTAGCC |
| 1_S288C_controls_B10 | CGTACTAG | GTCTGAGG |
| 1_S288C_controls_B11 | CGTACTAG | CGTAAGGA |
| 1_S288C_controls_B12 | CGTACTAG | CCACGCGT |
| 1_S288C_controls_C1 | AGGCAGAA | GCGATCTA |
| 1_S288C_controls_C2 | AGGCAGAA | ATAGAGAG |
| 1_S288C_controls_C3 | AGGCAGAA | AGAGGATA |
| 1_S288C_controls_C4 | AGGCAGAA | TCTACTCT |
| 1_S288C_controls_C5 | AGGCAGAA | CTCCTTAC |
| 1_S288C_controls_C6 | AGGCAGAA | TATGCAGT |
| 1_S288C_controls_C7 | AGGCAGAA | TACTCCTT |
| 1_S288C_controls_C8 | AGGCAGAA | AGGCTTAG |
| 1_S288C_controls_C9 | AGGCAGAA | GAGTAGCC |
| 1_S288C_controls_C10 | AGGCAGAA | GTCTGAGG |
| 1_S288C_controls_C11 | AGGCAGAA | CGTAAGGA |
| 1_S288C_controls_C12 | AGGCAGAA | CCACGCGT |
| 1_S288C_controls_D1 | TCCTGAGC | GCGATCTA |
| 1_S288C_controls_D2 | TCCTGAGC | ATAGAGAG |
| 1_S288C_controls_D3 | TCCTGAGC | AGAGGATA |
| 1_S288C_controls_D4 | TCCTGAGC | TCTACTCT |

| | | |
|---|---|---|
| 1_S288C_controls_D5 | TCCTGAGC | CTCCTTAC |
| 1_S288C_controls_D6 | TCCTGAGC | TATGCAGT |
| 1_S288C_controls_D7 | TCCTGAGC | TACTCCTT |
| 1_S288C_controls_D8 | TCCTGAGC | AGGCTTAG |
| 1_S288C_controls_D9 | TCCTGAGC | GAGTAGCC |
| 1_S288C_controls_D10 | TCCTGAGC | GTCTGAGG |
| 1_S288C_controls_D11 | TCCTGAGC | CGTAAGGA |
| 1_S288C_controls_D12 | TCCTGAGC | CCACGCGT |
| 1_S288C_controls_E1 | GGACTCCT | GCGATCTA |
| 1_S288C_controls_E2 | GGACTCCT | ATAGAGAG |
| 1_S288C_controls_E3 | GGACTCCT | AGAGGATA |
| 1_S288C_controls_E4 | GGACTCCT | TCTACTCT |
| 1_S288C_controls_E5 | GGACTCCT | CTCCTTAC |
| 1_S288C_controls_E6 | GGACTCCT | TATGCAGT |
| 1_S288C_controls_E7 | GGACTCCT | TACTCCTT |
| 1_S288C_controls_E8 | GGACTCCT | AGGCTTAG |
| 1_S288C_controls_E9 | GGACTCCT | GAGTAGCC |
| 1_S288C_controls_E10 | GGACTCCT | GTCTGAGG |
| 1_S288C_controls_E11 | GGACTCCT | CGTAAGGA |
| 1_S288C_controls_E12 | GGACTCCT | CCACGCGT |
| 1_S288C_controls_F1 | TAGGCATG | GCGATCTA |
| 1_S288C_controls_F2 | TAGGCATG | ATAGAGAG |
| 1_S288C_controls_F3 | TAGGCATG | AGAGGATA |
| 1_S288C_controls_F4 | TAGGCATG | TCTACTCT |
| 1_S288C_controls_F5 | TAGGCATG | CTCCTTAC |
| 1_S288C_controls_F6 | TAGGCATG | TATGCAGT |
| 1_S288C_controls_F7 | TAGGCATG | TACTCCTT |
| 1_S288C_controls_F8 | TAGGCATG | AGGCTTAG |
| 1_S288C_controls_F9 | TAGGCATG | GAGTAGCC |
| 1_S288C_controls_F10 | TAGGCATG | GTCTGAGG |
| 1_S288C_controls_F11 | TAGGCATG | CGTAAGGA |
| 1_S288C_controls_F12 | TAGGCATG | CCACGCGT |
| 1_S288C_controls_G1 | CTCTCTAC | GCGATCTA |
| 1_S288C_controls_G2 | CTCTCTAC | ATAGAGAG |
| 1_S288C_controls_G3 | CTCTCTAC | AGAGGATA |
| 1_S288C_controls_G4 | CTCTCTAC | TCTACTCT |
| 1_S288C_controls_G5 | CTCTCTAC | CTCCTTAC |
| 1_S288C_controls_G6 | CTCTCTAC | TATGCAGT |
| 1_S288C_controls_G7 | CTCTCTAC | TACTCCTT |
| 1_S288C_controls_G8 | CTCTCTAC | AGGCTTAG |
| 1_S288C_controls_G9 | CTCTCTAC | GAGTAGCC |
| 1_S288C_controls_G10 | CTCTCTAC | GTCTGAGG |
| 1_S288C_controls_G11 | CTCTCTAC | CGTAAGGA |

| | | |
|---|---|---|
| 1_S288C_controls_G12 | CTCTCTAC | CCACGCGT |
| 1_S288C_parental_H1 | CAGAGAGG | GCGATCTA |
| 1_S288C_parental_H2 | CAGAGAGG | ATAGAGAG |
| 1_S288C_parental_H3 | CAGAGAGG | AGAGGATA |
| 1_S288C_parental_H4 | CAGAGAGG | TCTACTCT |
| 1_S288C_parental_H5 | CAGAGAGG | CTCCTTAC |
| 1_S288C_parental_H6 | CAGAGAGG | TATGCAGT |
| 1_S288C_parental_H7 | CAGAGAGG | TACTCCTT |
| 1_S288C_parental_H8 | CAGAGAGG | AGGCTTAG |
| 1_S288C_controls_H9 | CAGAGAGG | GAGTAGCC |
| 1_S288C_controls_H10 | CAGAGAGG | GTCTGAGG |
| 1_S288C_controls_H11 | CAGAGAGG | CGTAAGGA |
| 1_S288C_controls_H12 | CAGAGAGG | CCACGCGT |
| 3_ETRED_controls_A1 | TAAGGCGA | GAATTCAG |
| 3_ETRED_controls_A2 | TAAGGCGA | CCGGTACG |
| 3_ETRED_controls_A3 | TAAGGCGA | CCGTCATC |
| 3_ETRED_controls_A4 | TAAGGCGA | CGTCTATA |
| 3_ETRED_controls_A5 | TAAGGCGA | TCAATGAC |
| 3_ETRED_controls_A6 | TAAGGCGA | AACGATGC |
| 3_ETRED_controls_A7 | TAAGGCGA | GTCAACCT |
| 3_ETRED_controls_A8 | TAAGGCGA | CAGTTTCA |
| 3_ETRED_controls_A9 | TAAGGCGA | TGTGATTG |
| 3_ETRED_controls_A10 | TAAGGCGA | TTGCATGT |
| 3_ETRED_controls_A11 | TAAGGCGA | GGCGCGAT |
| 3_ETRED_controls_A12 | TAAGGCGA | TTAACCGA |
| 3_ETRED_controls_B1 | CGTACTAG | GAATTCAG |
| 3_ETRED_controls_B2 | CGTACTAG | CCGGTACG |
| 3_ETRED_controls_B3 | CGTACTAG | CCGTCATC |
| 3_ETRED_controls_B4 | CGTACTAG | CGTCTATA |
| 3_ETRED_controls_B5 | CGTACTAG | TCAATGAC |
| 3_ETRED_controls_B6 | CGTACTAG | AACGATGC |
| 3_ETRED_controls_B7 | CGTACTAG | GTCAACCT |
| 3_ETRED_controls_B8 | CGTACTAG | CAGTTTCA |
| 3_ETRED_controls_B9 | CGTACTAG | TGTGATTG |
| 3_ETRED_controls_B10 | CGTACTAG | TTGCATGT |
| 3_ETRED_controls_B11 | CGTACTAG | GGCGCGAT |
| 3_ETRED_controls_B12 | CGTACTAG | TTAACCGA |
| 3_ETRED_controls_C1 | AGGCAGAA | GAATTCAG |
| 3_ETRED_controls_C2 | AGGCAGAA | CCGGTACG |
| 3_ETRED_controls_C3 | AGGCAGAA | CCGTCATC |
| 3_ETRED_controls_C4 | AGGCAGAA | CGTCTATA |
| 3_ETRED_controls_C5 | AGGCAGAA | TCAATGAC |
| 3_ETRED_controls_C6 | AGGCAGAA | AACGATGC |

| | | |
|---|---|---|
| 3_ETRED_controls_C7 | AGGCAGAA | GTCAACCT |
| 3_ETRED_controls_C8 | AGGCAGAA | CAGTTTCA |
| 3_ETRED_controls_C9 | AGGCAGAA | TGTGATTG |
| 3_ETRED_controls_C10 | AGGCAGAA | TTGCATGT |
| 3_ETRED_controls_C11 | AGGCAGAA | GGCGCGAT |
| 3_ETRED_controls_C12 | AGGCAGAA | TTAACCGA |
| 3_ETRED_controls_D1 | TCCTGAGC | GAATTCAG |
| 3_ETRED_controls_D2 | TCCTGAGC | CCGGTACG |
| 3_ETRED_controls_D3 | TCCTGAGC | CCGTCATC |
| 3_ETRED_controls_D4 | TCCTGAGC | CGTCTATA |
| 3_ETRED_controls_D5 | TCCTGAGC | TCAATGAC |
| 3_ETRED_controls_D6 | TCCTGAGC | AACGATGC |
| 3_ETRED_controls_D7 | TCCTGAGC | GTCAACCT |
| 3_ETRED_controls_D8 | TCCTGAGC | CAGTTTCA |
| 3_ETRED_controls_D9 | TCCTGAGC | TGTGATTG |
| 3_ETRED_controls_D10 | TCCTGAGC | TTGCATGT |
| 3_ETRED_controls_D11 | TCCTGAGC | GGCGCGAT |
| 3_ETRED_controls_D12 | TCCTGAGC | TTAACCGA |
| 3_ETRED_controls_E1 | GGACTCCT | GAATTCAG |
| 3_ETRED_controls_E2 | GGACTCCT | CCGGTACG |
| 3_ETRED_controls_E3 | GGACTCCT | CCGTCATC |
| 3_ETRED_controls_E4 | GGACTCCT | CGTCTATA |
| 3_ETRED_controls_E5 | GGACTCCT | TCAATGAC |
| 3_ETRED_controls_E6 | GGACTCCT | AACGATGC |
| 3_ETRED_controls_E7 | GGACTCCT | GTCAACCT |
| 3_ETRED_controls_E8 | GGACTCCT | CAGTTTCA |
| 3_ETRED_controls_E9 | GGACTCCT | TGTGATTG |
| 3_ETRED_controls_E10 | GGACTCCT | TTGCATGT |
| 3_ETRED_controls_E11 | GGACTCCT | GGCGCGAT |
| 3_ETRED_controls_E12 | GGACTCCT | TTAACCGA |
| 3_ETRED_parental_F1 | TAGGCATG | GAATTCAG |
| 3_ETRED_parental_F2 | TAGGCATG | CCGGTACG |
| 3_ETRED_parental_F3 | TAGGCATG | CCGTCATC |
| 3_ETRED_parental_F4 | TAGGCATG | CGTCTATA |
| 3_ETRED_parental_F5 | TAGGCATG | TCAATGAC |
| 3_ETRED_parental_F6 | TAGGCATG | AACGATGC |
| 3_ETRED_parental_F7 | TAGGCATG | GTCAACCT |
| 3_ETRED_parental_F8 | TAGGCATG | CAGTTTCA |
| 3_ETRED_controls_F9 | TAGGCATG | TGTGATTG |
| 3_ETRED_controls_F10 | TAGGCATG | TTGCATGT |
| 3_ETRED_controls_F11 | TAGGCATG | GGCGCGAT |
| 3_ETRED_controls_F12 | TAGGCATG | TTAACCGA |
| 3_ETRED_controls_G1 | CTCTCTAC | GAATTCAG |

| | | |
|---|---|---|
| 3_ETRED_controls_G2 | CTCTCTAC | CCGGTACG |
| 3_ETRED_controls_G3 | CTCTCTAC | CCGTCATC |
| 3_ETRED_controls_G4 | CTCTCTAC | CGTCTATA |
| 3_ETRED_controls_G5 | CTCTCTAC | TCAATGAC |
| 3_ETRED_controls_G6 | CTCTCTAC | AACGATGC |
| 3_ETRED_controls_G7 | CTCTCTAC | GTCAACCT |
| 3_ETRED_controls_G8 | CTCTCTAC | CAGTTTCA |
| 3_ETRED_controls_G9 | CTCTCTAC | TGTGATTG |
| 3_ETRED_controls_G10 | CTCTCTAC | TTGCATGT |
| 3_ETRED_controls_G11 | CTCTCTAC | GGCGCGAT |
| 3_ETRED_controls_G12 | CTCTCTAC | TTAACCGA |
| 3_ETRED_controls_H1 | CAGAGAGG | GAATTCAG |
| 3_ETRED_controls_H2 | CAGAGAGG | CCGGTACG |
| 3_ETRED_controls_H3 | CAGAGAGG | CCGTCATC |
| 3_ETRED_controls_H4 | CAGAGAGG | CGTCTATA |
| 3_ETRED_controls_H5 | CAGAGAGG | TCAATGAC |
| 3_ETRED_controls_H6 | CAGAGAGG | AACGATGC |
| 3_ETRED_controls_H7 | CAGAGAGG | GTCAACCT |
| 3_ETRED_controls_H8 | CAGAGAGG | CAGTTTCA |
| 3_ETRED_controls_H9 | CAGAGAGG | TGTGATTG |
| 3_ETRED_controls_H10 | CAGAGAGG | TTGCATGT |
| 3_ETRED_controls_H11 | CAGAGAGG | GGCGCGAT |
| 3_ETRED_controls_H12 | CAGAGAGG | TTAACCGA |
| 4_S288C_G8EL1_A1 | GCTACGCT | GCGATCTA |
| 4_S288C_G8EL1_A2 | GCTACGCT | ATAGAGAG |
| 4_S288C_G8EL1_A3 | GCTACGCT | AGAGGATA |
| 4_S288C_G8EL1_A4 | GCTACGCT | TCTACTCT |
| 4_S288C_G8EL2_A5 | GCTACGCT | CTCCTTAC |
| 4_S288C_G8EL2_A6 | GCTACGCT | TATGCAGT |
| 4_S288C_G8EL2_A7 | GCTACGCT | TACTCCTT |
| 4_S288C_G8EL2_A8 | GCTACGCT | AGGCTTAG |
| 4_S288C_G8EL3_A9 | GCTACGCT | GAGTAGCC |
| 4_S288C_G8EL3_A10 | GCTACGCT | GTCTGAGG |
| 4_S288C_G8EL3_A11 | GCTACGCT | CGTAAGGA |
| 4_S288C_G8EL3_A12 | GCTACGCT | CCACGCGT |
| 4_S288C_G8EL1_B1 | CGAGGCTG | GCGATCTA |
| 4_S288C_G8EL1_B2 | CGAGGCTG | ATAGAGAG |
| 4_S288C_G8EL1_B3 | CGAGGCTG | AGAGGATA |
| 4_S288C_G8EL1_B4 | CGAGGCTG | TCTACTCT |
| 4_S288C_G8EL2_B5 | CGAGGCTG | CTCCTTAC |
| 4_S288C_G8EL2_B6 | CGAGGCTG | TATGCAGT |
| 4_S288C_G8EL2_B7 | CGAGGCTG | TACTCCTT |
| 4_S288C_G8EL2_B8 | CGAGGCTG | AGGCTTAG |

| | | |
|---|---|---|
| 4_S288C_G8EL3_B9 | CGAGGCTG | GAGTAGCC |
| 4_S288C_G8EL3_B10 | CGAGGCTG | GTCTGAGG |
| 4_S288C_G8EL3_B11 | CGAGGCTG | CGTAAGGA |
| 4_S288C_G8EL3_B12 | CGAGGCTG | CCACGCGT |
| 4_S288C_G8EL1_C1 | AAGAGGCA | GCGATCTA |
| 4_S288C_G8EL1_C2 | AAGAGGCA | ATAGAGAG |
| 4_S288C_G8EL1_C3 | AAGAGGCA | AGAGGATA |
| 4_S288C_G8EL1_C4 | AAGAGGCA | TCTACTCT |
| 4_S288C_G8EL2_C5 | AAGAGGCA | CTCCTTAC |
| 4_S288C_G8EL2_C6 | AAGAGGCA | TATGCAGT |
| 4_S288C_G8EL2_C7 | AAGAGGCA | TACTCCTT |
| 4_S288C_G8EL2_C8 | AAGAGGCA | AGGCTTAG |
| 4_S288C_G8EL3_C9 | AAGAGGCA | GAGTAGCC |
| 4_S288C_G8EL3_C10 | AAGAGGCA | GTCTGAGG |
| 4_S288C_G8EL3_C11 | AAGAGGCA | CGTAAGGA |
| 4_S288C_G8EL3_C12 | AAGAGGCA | CCACGCGT |
| 4_S288C_G8EL1_D1 | GTAGAGGA | GCGATCTA |
| 4_S288C_G8EL1_D2 | GTAGAGGA | ATAGAGAG |
| 4_S288C_G8EL1_D3 | GTAGAGGA | AGAGGATA |
| 4_S288C_G8EL1_D4 | GTAGAGGA | TCTACTCT |
| 4_S288C_G8EL2_D5 | GTAGAGGA | CTCCTTAC |
| 4_S288C_G8EL2_D6 | GTAGAGGA | TATGCAGT |
| 4_S288C_G8EL2_D7 | GTAGAGGA | TACTCCTT |
| 4_S288C_G8EL2_D8 | GTAGAGGA | AGGCTTAG |
| 4_S288C_G8EL3_D9 | GTAGAGGA | GAGTAGCC |
| 4_S288C_G8EL3_D10 | GTAGAGGA | GTCTGAGG |
| 4_S288C_G8EL3_D11 | GTAGAGGA | CGTAAGGA |
| 4_S288C_G8EL3_D12 | GTAGAGGA | CCACGCGT |
| 4_S288C_G8EL1_E1 | ATTGTAAT | GCGATCTA |
| 4_S288C_G8EL1_E2 | ATTGTAAT | ATAGAGAG |
| 4_S288C_G8EL1_E3 | ATTGTAAT | AGAGGATA |
| 4_S288C_G8EL1_E4 | ATTGTAAT | TCTACTCT |
| 4_S288C_G8EL2_E5 | ATTGTAAT | CTCCTTAC |
| 4_S288C_G8EL2_E6 | ATTGTAAT | TATGCAGT |
| 4_S288C_G8EL2_E7 | ATTGTAAT | TACTCCTT |
| 4_S288C_G8EL2_E8 | ATTGTAAT | AGGCTTAG |
| 4_S288C_G8EL3_E9 | ATTGTAAT | GAGTAGCC |
| 4_S288C_G8EL3_E10 | ATTGTAAT | GTCTGAGG |
| 4_S288C_G8EL3_E11 | ATTGTAAT | CGTAAGGA |
| 4_S288C_G8EL3_E12 | ATTGTAAT | CCACGCGT |
| 4_S288C_G8EL1_F1 | GATCATTC | GCGATCTA |
| 4_S288C_G8EL1_F2 | GATCATTC | ATAGAGAG |
| 4_S288C_G8EL1_F3 | GATCATTC | AGAGGATA |

| | | |
|---|---|---|
| 4_S288C_G8EL1_F4 | GATCATTC | TCTACTCT |
| 4_S288C_G8EL2_F5 | GATCATTC | CTCCTTAC |
| 4_S288C_G8EL2_F6 | GATCATTC | TATGCAGT |
| 4_S288C_G8EL2_F7 | GATCATTC | TACTCCTT |
| 4_S288C_G8EL2_F8 | GATCATTC | AGGCTTAG |
| 4_S288C_G8EL3_F9 | GATCATTC | GAGTAGCC |
| 4_S288C_G8EL3_F10 | GATCATTC | GTCTGAGG |
| 4_S288C_G8EL3_F11 | GATCATTC | CGTAAGGA |
| 4_S288C_G8EL3_F12 | GATCATTC | CCACGCGT |
| 4_S288C_G8EL1_G1 | ACCGATCG | GCGATCTA |
| 4_S288C_G8EL1_G2 | ACCGATCG | ATAGAGAG |
| 4_S288C_G8EL1_G3 | ACCGATCG | AGAGGATA |
| 4_S288C_G8EL1_G4 | ACCGATCG | TCTACTCT |
| 4_S288C_G8EL2_G5 | ACCGATCG | CTCCTTAC |
| 4_S288C_G8EL2_G6 | ACCGATCG | TATGCAGT |
| 4_S288C_G8EL2_G7 | ACCGATCG | TACTCCTT |
| 4_S288C_G8EL2_G8 | ACCGATCG | AGGCTTAG |
| 4_S288C_G8EL3_G9 | ACCGATCG | GAGTAGCC |
| 4_S288C_G8EL3_G10 | ACCGATCG | GTCTGAGG |
| 4_S288C_G8EL3_G11 | ACCGATCG | CGTAAGGA |
| 4_S288C_G8EL3_G12 | ACCGATCG | CCACGCGT |
| 4_S288C_G8EL1_H1 | CCGTTATT | GCGATCTA |
| 4_S288C_G8EL1_H2 | CCGTTATT | ATAGAGAG |
| 4_S288C_G8EL1_H3 | CCGTTATT | AGAGGATA |
| 4_S288C_G8EL1_H4 | CCGTTATT | TCTACTCT |
| 4_S288C_G8EL2_H5 | CCGTTATT | CTCCTTAC |
| 4_S288C_G8EL2_H6 | CCGTTATT | TATGCAGT |
| 4_S288C_G8EL2_H7 | CCGTTATT | TACTCCTT |
| 4_S288C_G8EL2_H8 | CCGTTATT | AGGCTTAG |
| 4_S288C_G8EL3_H9 | CCGTTATT | GAGTAGCC |
| 4_S288C_G8EL3_H10 | CCGTTATT | GTCTGAGG |
| 4_S288C_G8EL3_H11 | CCGTTATT | CGTAAGGA |
| 4_S288C_G8EL3_H12 | CCGTTATT | CCACGCGT |
| 6_ETRED_G8EL1_A1 | GCTACGCT | GAATTCAG |
| 6_ETRED_G8EL1_A2 | GCTACGCT | CCGGTACG |
| 6_ETRED_G8EL1_A3 | GCTACGCT | CCGTCATC |
| 6_ETRED_G8EL1_A4 | GCTACGCT | CGTCTATA |
| 6_ETRED_G8EL2_A5 | GCTACGCT | TCAATGAC |
| 6_ETRED_G8EL2_A6 | GCTACGCT | AACGATGC |
| 6_ETRED_G8EL2_A7 | GCTACGCT | GTCAACCT |
| 6_ETRED_G8EL2_A8 | GCTACGCT | CAGTTTCA |
| 6_ETRED_G8EL3_A9 | GCTACGCT | TGTGATTG |
| 6_ETRED_G8EL3_A10 | GCTACGCT | TTGCATGT |

| | | |
|---|---|---|
| 6_ETRED_G8EL3_A11 | GCTACGCT | GGCGCGAT |
| 6_ETRED_G8EL3_A12 | GCTACGCT | TTAACCGA |
| 6_ETRED_G8EL1_B1 | CGAGGCTG | GAATTCAG |
| 6_ETRED_G8EL1_B2 | CGAGGCTG | CCGGTACG |
| 6_ETRED_G8EL1_B3 | CGAGGCTG | CCGTCATC |
| 6_ETRED_G8EL1_B4 | CGAGGCTG | CGTCTATA |
| 6_ETRED_G8EL2_B5 | CGAGGCTG | TCAATGAC |
| 6_ETRED_G8EL2_B6 | CGAGGCTG | AACGATGC |
| 6_ETRED_G8EL2_B7 | CGAGGCTG | GTCAACCT |
| 6_ETRED_G8EL2_B8 | CGAGGCTG | CAGTTTCA |
| 6_ETRED_G8EL3_B9 | CGAGGCTG | TGTGATTG |
| 6_ETRED_G8EL3_B10 | CGAGGCTG | TTGCATGT |
| 6_ETRED_G8EL3_B11 | CGAGGCTG | GGCGCGAT |
| 6_ETRED_G8EL3_B12 | CGAGGCTG | TTAACCGA |
| 6_ETRED_G8EL1_C1 | AAGAGGCA | GAATTCAG |
| 6_ETRED_G8EL1_C2 | AAGAGGCA | CCGGTACG |
| 6_ETRED_G8EL1_C3 | AAGAGGCA | CCGTCATC |
| 6_ETRED_G8EL1_C4 | AAGAGGCA | CGTCTATA |
| 6_ETRED_G8EL2_C5 | AAGAGGCA | TCAATGAC |
| 6_ETRED_G8EL2_C6 | AAGAGGCA | AACGATGC |
| 6_ETRED_G8EL2_C7 | AAGAGGCA | GTCAACCT |
| 6_ETRED_G8EL2_C8 | AAGAGGCA | CAGTTTCA |
| 6_ETRED_G8EL3_C9 | AAGAGGCA | TGTGATTG |
| 6_ETRED_G8EL3_C10 | AAGAGGCA | TTGCATGT |
| 6_ETRED_G8EL3_C11 | AAGAGGCA | GGCGCGAT |
| 6_ETRED_G8EL3_C12 | AAGAGGCA | TTAACCGA |
| 6_ETRED_G8EL1_D1 | GTAGAGGA | GAATTCAG |
| 6_ETRED_G8EL1_D2 | GTAGAGGA | CCGGTACG |
| 6_ETRED_G8EL1_D3 | GTAGAGGA | CCGTCATC |
| 6_ETRED_G8EL1_D4 | GTAGAGGA | CGTCTATA |
| 6_ETRED_G8EL2_D5 | GTAGAGGA | TCAATGAC |
| 6_ETRED_G8EL2_D6 | GTAGAGGA | AACGATGC |
| 6_ETRED_G8EL2_D7 | GTAGAGGA | GTCAACCT |
| 6_ETRED_G8EL2_D8 | GTAGAGGA | CAGTTTCA |
| 6_ETRED_G8EL3_D9 | GTAGAGGA | TGTGATTG |
| 6_ETRED_G8EL3_D10 | GTAGAGGA | TTGCATGT |
| 6_ETRED_G8EL3_D11 | GTAGAGGA | GGCGCGAT |
| 6_ETRED_G8EL3_D12 | GTAGAGGA | TTAACCGA |
| 6_ETRED_G8EL1_E1 | ATTGTAAT | GAATTCAG |
| 6_ETRED_G8EL1_E2 | ATTGTAAT | CCGGTACG |
| 6_ETRED_G8EL1_E3 | ATTGTAAT | CCGTCATC |
| 6_ETRED_G8EL1_E4 | ATTGTAAT | CGTCTATA |
| 6_ETRED_G8EL2_E5 | ATTGTAAT | TCAATGAC |

| | | |
|---|---|---|
| 6_ETRED_G8EL2_E6 | ATTGTAAT | AACGATGC |
| 6_ETRED_G8EL2_E7 | ATTGTAAT | GTCAACCT |
| 6_ETRED_G8EL2_E8 | ATTGTAAT | CAGTTTCA |
| 6_ETRED_G8EL3_E9 | ATTGTAAT | TGTGATTG |
| 6_ETRED_G8EL3_E10 | ATTGTAAT | TTGCATGT |
| 6_ETRED_G8EL3_E11 | ATTGTAAT | GGCGCGAT |
| 6_ETRED_G8EL3_E12 | ATTGTAAT | TTAACCGA |
| 6_ETRED_G8EL1_F1 | GATCATTC | GAATTCAG |
| 6_ETRED_G8EL1_F2 | GATCATTC | CCGGTACG |
| 6_ETRED_G8EL1_F3 | GATCATTC | CCGTCATC |
| 6_ETRED_G8EL1_F4 | GATCATTC | CGTCTATA |
| 6_ETRED_G8EL2_F5 | GATCATTC | TCAATGAC |
| 6_ETRED_G8EL2_F6 | GATCATTC | AACGATGC |
| 6_ETRED_G8EL2_F7 | GATCATTC | GTCAACCT |
| 6_ETRED_G8EL2_F8 | GATCATTC | CAGTTTCA |
| 6_ETRED_G8EL3_F9 | GATCATTC | TGTGATTG |
| 6_ETRED_G8EL3_F10 | GATCATTC | TTGCATGT |
| 6_ETRED_G8EL3_F11 | GATCATTC | GGCGCGAT |
| 6_ETRED_G8EL3_F12 | GATCATTC | TTAACCGA |
| 6_ETRED_G8EL1_G1 | ACCGATCG | GAATTCAG |
| 6_ETRED_G8EL1_G2 | ACCGATCG | CCGGTACG |
| 6_ETRED_G8EL1_G3 | ACCGATCG | CCGTCATC |
| 6_ETRED_G8EL1_G4 | ACCGATCG | CGTCTATA |
| 6_ETRED_G8EL2_G5 | ACCGATCG | TCAATGAC |
| 6_ETRED_G8EL2_G6 | ACCGATCG | AACGATGC |
| 6_ETRED_G8EL2_G7 | ACCGATCG | GTCAACCT |
| 6_ETRED_G8EL2_G8 | ACCGATCG | CAGTTTCA |
| 6_ETRED_G8EL3_G9 | ACCGATCG | TGTGATTG |
| 6_ETRED_G8EL3_G10 | ACCGATCG | TTGCATGT |
| 6_ETRED_G8EL3_G11 | ACCGATCG | GGCGCGAT |
| 6_ETRED_G8EL3_G12 | ACCGATCG | TTAACCGA |
| 6_ETRED_G8EL1_H1 | CCGTTATT | GAATTCAG |
| 6_ETRED_G8EL1_H2 | CCGTTATT | CCGGTACG |
| 6_ETRED_G8EL1_H3 | CCGTTATT | CCGTCATC |
| 6_ETRED_G8EL1_H4 | CCGTTATT | CGTCTATA |
| 6_ETRED_G8EL2_H5 | CCGTTATT | TCAATGAC |
| 6_ETRED_G8EL2_H6 | CCGTTATT | AACGATGC |
| 6_ETRED_G8EL2_H7 | CCGTTATT | GTCAACCT |
| 6_ETRED_G8EL2_H8 | CCGTTATT | CAGTTTCA |
| 6_ETRED_G8EL3_H9 | CCGTTATT | TGTGATTG |
| 6_ETRED_G8EL3_H10 | CCGTTATT | TTGCATGT |
| 6_ETRED_G8EL3_H11 | CCGTTATT | GGCGCGAT |
| 6_ETRED_G8EL3_H12 | CCGTTATT | TTAACCGA |

| | | |
|---|---|---|
| 7_S288C_G80EL1_A1 | TTCTTCTA | GCGATCTA |
| 7_S288C_G80EL1_A2 | TTCTTCTA | ATAGAGAG |
| 7_S288C_G80EL1_A3 | TTCTTCTA | AGAGGATA |
| 7_S288C_G80EL1_A4 | TTCTTCTA | TCTACTCT |
| 7_S288C_G80EL2_A5 | TTCTTCTA | CTCCTTAC |
| 7_S288C_G80EL2_A6 | TTCTTCTA | TATGCAGT |
| 7_S288C_G80EL2_A7 | TTCTTCTA | TACTCCTT |
| 7_S288C_G80EL2_A8 | TTCTTCTA | AGGCTTAG |
| 7_S288C_G80EL3_A9 | TTCTTCTA | GAGTAGCC |
| 7_S288C_G80EL3_A10 | TTCTTCTA | GTCTGAGG |
| 7_S288C_G80EL3_A11 | TTCTTCTA | CGTAAGGA |
| 7_S288C_G80EL3_A12 | TTCTTCTA | CCACGCGT |
| 7_S288C_G80EL1_B1 | TACCTGAC | GCGATCTA |
| 7_S288C_G80EL1_B2 | TACCTGAC | ATAGAGAG |
| 7_S288C_G80EL1_B3 | TACCTGAC | AGAGGATA |
| 7_S288C_G80EL1_B4 | TACCTGAC | TCTACTCT |
| 7_S288C_G80EL2_B5 | TACCTGAC | CTCCTTAC |
| 7_S288C_G80EL2_B6 | TACCTGAC | TATGCAGT |
| 7_S288C_G80EL2_B7 | TACCTGAC | TACTCCTT |
| 7_S288C_G80EL2_B8 | TACCTGAC | AGGCTTAG |
| 7_S288C_G80EL3_B9 | TACCTGAC | GAGTAGCC |
| 7_S288C_G80EL3_B10 | TACCTGAC | GTCTGAGG |
| 7_S288C_G80EL3_B11 | TACCTGAC | CGTAAGGA |
| 7_S288C_G80EL3_B12 | TACCTGAC | CCACGCGT |
| 7_S288C_G80EL1_C1 | AGGACCGC | GCGATCTA |
| 7_S288C_G80EL1_C2 | AGGACCGC | ATAGAGAG |
| 7_S288C_G80EL1_C3 | AGGACCGC | AGAGGATA |
| 7_S288C_G80EL1_C4 | AGGACCGC | TCTACTCT |
| 7_S288C_G80EL2_C5 | AGGACCGC | CTCCTTAC |
| 7_S288C_G80EL2_C6 | AGGACCGC | TATGCAGT |
| 7_S288C_G80EL2_C7 | AGGACCGC | TACTCCTT |
| 7_S288C_G80EL2_C8 | AGGACCGC | AGGCTTAG |
| 7_S288C_G80EL3_C9 | AGGACCGC | GAGTAGCC |
| 7_S288C_G80EL3_C10 | AGGACCGC | GTCTGAGG |
| 7_S288C_G80EL3_C11 | AGGACCGC | CGTAAGGA |
| 7_S288C_G80EL3_C12 | AGGACCGC | CCACGCGT |
| 7_S288C_G80EL1_D1 | GTCCGATT | GCGATCTA |
| 7_S288C_G80EL1_D2 | GTCCGATT | ATAGAGAG |
| 7_S288C_G80EL1_D3 | GTCCGATT | AGAGGATA |
| 7_S288C_G80EL1_D4 | GTCCGATT | TCTACTCT |
| 7_S288C_G80EL2_D5 | GTCCGATT | CTCCTTAC |
| 7_S288C_G80EL2_D6 | GTCCGATT | TATGCAGT |
| 7_S288C_G80EL2_D7 | GTCCGATT | TACTCCTT |

| | | |
|---|---|---|
| 7_S288C_G80EL2_D8 | GTCCGATT | AGGCTTAG |
| 7_S288C_G80EL3_D9 | GTCCGATT | GAGTAGCC |
| 7_S288C_G80EL3_D10 | GTCCGATT | GTCTGAGG |
| 7_S288C_G80EL3_D11 | GTCCGATT | CGTAAGGA |
| 7_S288C_G80EL3_D12 | GTCCGATT | CCACGCGT |
| 7_S288C_G80EL1_E1 | CACGAGTT | GCGATCTA |
| 7_S288C_G80EL1_E2 | CACGAGTT | ATAGAGAG |
| 7_S288C_G80EL1_E3 | CACGAGTT | AGAGGATA |
| 7_S288C_G80EL1_E4 | CACGAGTT | TCTACTCT |
| 7_S288C_G80EL2_E5 | CACGAGTT | CTCCTTAC |
| 7_S288C_G80EL2_E6 | CACGAGTT | TATGCAGT |
| 7_S288C_G80EL2_E7 | CACGAGTT | TACTCCTT |
| 7_S288C_G80EL2_E8 | CACGAGTT | AGGCTTAG |
| 7_S288C_G80EL3_E9 | CACGAGTT | GAGTAGCC |
| 7_S288C_G80EL3_E10 | CACGAGTT | GTCTGAGG |
| 7_S288C_G80EL3_E11 | CACGAGTT | CGTAAGGA |
| 7_S288C_G80EL3_E12 | CACGAGTT | CCACGCGT |
| 7_S288C_G80EL1_F1 | CCACGGCC | GCGATCTA |
| 7_S288C_G80EL1_F2 | CCACGGCC | ATAGAGAG |
| 7_S288C_G80EL1_F3 | CCACGGCC | AGAGGATA |
| 7_S288C_G80EL1_F4 | CCACGGCC | TCTACTCT |
| 7_S288C_G80EL2_F5 | CCACGGCC | CTCCTTAC |
| 7_S288C_G80EL2_F6 | CCACGGCC | TATGCAGT |
| 7_S288C_G80EL2_F7 | CCACGGCC | TACTCCTT |
| 7_S288C_G80EL2_F8 | CCACGGCC | AGGCTTAG |
| 7_S288C_G80EL3_F9 | CCACGGCC | GAGTAGCC |
| 7_S288C_G80EL3_F10 | CCACGGCC | GTCTGAGG |
| 7_S288C_G80EL3_F11 | CCACGGCC | CGTAAGGA |
| 7_S288C_G80EL3_F12 | CCACGGCC | CCACGCGT |
| 7_S288C_G80EL1_G1 | ACATGTAA | GCGATCTA |
| 7_S288C_G80EL1_G2 | ACATGTAA | ATAGAGAG |
| 7_S288C_G80EL1_G3 | ACATGTAA | AGAGGATA |
| 7_S288C_G80EL1_G4 | ACATGTAA | TCTACTCT |
| 7_S288C_G80EL2_G5 | ACATGTAA | CTCCTTAC |
| 7_S288C_G80EL2_G6 | ACATGTAA | TATGCAGT |
| 7_S288C_G80EL2_G7 | ACATGTAA | TACTCCTT |
| 7_S288C_G80EL2_G8 | ACATGTAA | AGGCTTAG |
| 7_S288C_G80EL3_G9 | ACATGTAA | GAGTAGCC |
| 7_S288C_G80EL3_G10 | ACATGTAA | GTCTGAGG |
| 7_S288C_G80EL3_G11 | ACATGTAA | CGTAAGGA |
| 7_S288C_G80EL3_G12 | ACATGTAA | CCACGCGT |
| 7_S288C_G80EL1_H1 | TGTTAACT | GCGATCTA |
| 7_S288C_G80EL1_H2 | TGTTAACT | ATAGAGAG |

| | | |
|---|---|---|
| 7_S288C_G80EL1_H3 | TGTTAACT | AGAGGATA |
| 7_S288C_G80EL1_H4 | TGTTAACT | TCTACTCT |
| 7_S288C_G80EL2_H5 | TGTTAACT | CTCCTTAC |
| 7_S288C_G80EL2_H6 | TGTTAACT | TATGCAGT |
| 7_S288C_G80EL2_H7 | TGTTAACT | TACTCCTT |
| 7_S288C_G80EL2_H8 | TGTTAACT | AGGCTTAG |
| 7_S288C_G80EL3_H9 | TGTTAACT | GAGTAGCC |
| 7_S288C_G80EL3_H10 | TGTTAACT | GTCTGAGG |
| 7_S288C_G80EL3_H11 | TGTTAACT | CGTAAGGA |
| 7_S288C_G80EL3_H12 | TGTTAACT | CCACGCGT |
| 9_ETRED_G80EL1_A1 | TTCTTCTA | GAATTCAG |
| 9_ETRED_G80EL1_A2 | TTCTTCTA | CCGGTACG |
| 9_ETRED_G80EL1_A3 | TTCTTCTA | CCGTCATC |
| 9_ETRED_G80EL1_A4 | TTCTTCTA | CGTCTATA |
| 9_ETRED_G80EL2_A5 | TTCTTCTA | TCAATGAC |
| 9_ETRED_G80EL2_A6 | TTCTTCTA | AACGATGC |
| 9_ETRED_G80EL2_A7 | TTCTTCTA | GTCAACCT |
| 9_ETRED_G80EL2_A8 | TTCTTCTA | CAGTTTCA |
| 9_ETRED_G80EL3_A9 | TTCTTCTA | TGTGATTG |
| 9_ETRED_G80EL3_A10 | TTCTTCTA | TTGCATGT |
| 9_ETRED_G80EL3_A11 | TTCTTCTA | GGCGCGAT |
| 9_ETRED_G80EL3_A12 | TTCTTCTA | TTAACCGA |
| 9_ETRED_G80EL1_B1 | TACCTGAC | GAATTCAG |
| 9_ETRED_G80EL1_B2 | TACCTGAC | CCGGTACG |
| 9_ETRED_G80EL1_B3 | TACCTGAC | CCGTCATC |
| 9_ETRED_G80EL1_B4 | TACCTGAC | CGTCTATA |
| 9_ETRED_G80EL2_B5 | TACCTGAC | TCAATGAC |
| 9_ETRED_G80EL2_B6 | TACCTGAC | AACGATGC |
| 9_ETRED_G80EL2_B7 | TACCTGAC | GTCAACCT |
| 9_ETRED_G80EL2_B8 | TACCTGAC | CAGTTTCA |
| 9_ETRED_G80EL3_B9 | TACCTGAC | TGTGATTG |
| 9_ETRED_G80EL3_B10 | TACCTGAC | TTGCATGT |
| 9_ETRED_G80EL3_B11 | TACCTGAC | GGCGCGAT |
| 9_ETRED_G80EL3_B12 | TACCTGAC | TTAACCGA |
| 9_ETRED_G80EL1_C1 | AGGACCGC | GAATTCAG |
| 9_ETRED_G80EL1_C2 | AGGACCGC | CCGGTACG |
| 9_ETRED_G80EL1_C3 | AGGACCGC | CCGTCATC |
| 9_ETRED_G80EL1_C4 | AGGACCGC | CGTCTATA |
| 9_ETRED_G80EL2_C5 | AGGACCGC | TCAATGAC |
| 9_ETRED_G80EL2_C6 | AGGACCGC | AACGATGC |
| 9_ETRED_G80EL2_C7 | AGGACCGC | GTCAACCT |
| 9_ETRED_G80EL2_C8 | AGGACCGC | CAGTTTCA |
| 9_ETRED_G80EL3_C9 | AGGACCGC | TGTGATTG |

| | | |
|---|---|---|
| 9_ETRED_G80EL3_C10 | AGGACCGC | TTGCATGT |
| 9_ETRED_G80EL3_C11 | AGGACCGC | GGCGCGAT |
| 9_ETRED_G80EL3_C12 | AGGACCGC | TTAACCGA |
| 9_ETRED_G80EL1_D1 | GTCCGATT | GAATTCAG |
| 9_ETRED_G80EL1_D2 | GTCCGATT | CCGGTACG |
| 9_ETRED_G80EL1_D3 | GTCCGATT | CCGTCATC |
| 9_ETRED_G80EL1_D4 | GTCCGATT | CGTCTATA |
| 9_ETRED_G80EL2_D5 | GTCCGATT | TCAATGAC |
| 9_ETRED_G80EL2_D6 | GTCCGATT | AACGATGC |
| 9_ETRED_G80EL2_D7 | GTCCGATT | GTCAACCT |
| 9_ETRED_G80EL2_D8 | GTCCGATT | CAGTTTCA |
| 9_ETRED_G80EL3_D9 | GTCCGATT | TGTGATTG |
| 9_ETRED_G80EL3_D10 | GTCCGATT | TTGCATGT |
| 9_ETRED_G80EL3_D11 | GTCCGATT | GGCGCGAT |
| 9_ETRED_G80EL3_D12 | GTCCGATT | TTAACCGA |
| 9_ETRED_G80EL1_E1 | CACGAGTT | GAATTCAG |
| 9_ETRED_G80EL1_E2 | CACGAGTT | CCGGTACG |
| 9_ETRED_G80EL1_E3 | CACGAGTT | CCGTCATC |
| 9_ETRED_G80EL1_E4 | CACGAGTT | CGTCTATA |
| 9_ETRED_G80EL2_E5 | CACGAGTT | TCAATGAC |
| 9_ETRED_G80EL2_E6 | CACGAGTT | AACGATGC |
| 9_ETRED_G80EL2_E7 | CACGAGTT | GTCAACCT |
| 9_ETRED_G80EL2_E8 | CACGAGTT | CAGTTTCA |
| 9_ETRED_G80EL3_E9 | CACGAGTT | TGTGATTG |
| 9_ETRED_G80EL3_E10 | CACGAGTT | TTGCATGT |
| 9_ETRED_G80EL3_E11 | CACGAGTT | GGCGCGAT |
| 9_ETRED_G80EL3_E12 | CACGAGTT | TTAACCGA |
| 9_ETRED_G80EL1_F1 | CCACGGCC | GAATTCAG |
| 9_ETRED_G80EL1_F2 | CCACGGCC | CCGGTACG |
| 9_ETRED_G80EL1_F3 | CCACGGCC | CCGTCATC |
| 9_ETRED_G80EL1_F4 | CCACGGCC | CGTCTATA |
| 9_ETRED_G80EL2_F5 | CCACGGCC | TCAATGAC |
| 9_ETRED_G80EL2_F6 | CCACGGCC | AACGATGC |
| 9_ETRED_G80EL2_F7 | CCACGGCC | GTCAACCT |
| 9_ETRED_G80EL2_F8 | CCACGGCC | CAGTTTCA |
| 9_ETRED_G80EL3_F9 | CCACGGCC | TGTGATTG |
| 9_ETRED_G80EL3_F10 | CCACGGCC | TTGCATGT |
| 9_ETRED_G80EL3_F11 | CCACGGCC | GGCGCGAT |
| 9_ETRED_G80EL3_F12 | CCACGGCC | TTAACCGA |
| 9_ETRED_G80EL1_G1 | ACATGTAA | GAATTCAG |
| 9_ETRED_G80EL1_G2 | ACATGTAA | CCGGTACG |
| 9_ETRED_G80EL1_G3 | ACATGTAA | CCGTCATC |
| 9_ETRED_G80EL1_G4 | ACATGTAA | CGTCTATA |

| | | |
|---|---|---|
| 9_ETRED_G80EL2_G5 | ACATGTAA | TCAATGAC |
| 9_ETRED_G80EL2_G6 | ACATGTAA | AACGATGC |
| 9_ETRED_G80EL2_G7 | ACATGTAA | GTCAACCT |
| 9_ETRED_G80EL2_G8 | ACATGTAA | CAGTTTCA |
| 9_ETRED_G80EL3_G9 | ACATGTAA | TGTGATTG |
| 9_ETRED_G80EL3_G10 | ACATGTAA | TTGCATGT |
| 9_ETRED_G80EL3_G11 | ACATGTAA | GGCGCGAT |
| 9_ETRED_G80EL3_G12 | ACATGTAA | TTAACCGA |
| 9_ETRED_G80EL1_H1 | TGTTAACT | GAATTCAG |
| 9_ETRED_G80EL1_H2 | TGTTAACT | CCGGTACG |
| 9_ETRED_G80EL1_H3 | TGTTAACT | CCGTCATC |
| 9_ETRED_G80EL1_H4 | TGTTAACT | CGTCTATA |
| 9_ETRED_G80EL2_H5 | TGTTAACT | TCAATGAC |
| 9_ETRED_G80EL2_H6 | TGTTAACT | AACGATGC |
| 9_ETRED_G80EL2_H7 | TGTTAACT | GTCAACCT |
| 9_ETRED_G80EL2_H8 | TGTTAACT | CAGTTTCA |
| 9_ETRED_G80EL3_H9 | TGTTAACT | TGTGATTG |
| 9_ETRED_G80EL3_H10 | TGTTAACT | TTGCATGT |
| 9_ETRED_G80EL3_H11 | TGTTAACT | GGCGCGAT |
| 9_ETRED_G80EL3_H12 | TGTTAACT | TTAACCGA |
| 10_S288C_controls_A1 | ATGGAAGC | GCGATCTA |
| 10_S288C_controls_A2 | ATGGAAGC | ATAGAGAG |
| 10_S288C_controls_A3 | ATGGAAGC | AGAGGATA |
| 10_S288C_controls_A4 | ATGGAAGC | TCTACTCT |
| 10_S288C_controls_A5 | ATGGAAGC | CTCCTTAC |
| 10_S288C_controls_A6 | ATGGAAGC | TATGCAGT |
| 10_S288C_controls_B1 | ATGTGGTG | GCGATCTA |
| 10_S288C_controls_B2 | ATGTGGTG | ATAGAGAG |
| 10_S288C_controls_B3 | ATGTGGTG | AGAGGATA |
| 10_S288C_controls_B4 | ATGTGGTG | TCTACTCT |
| 10_S288C_controls_B5 | ATGTGGTG | CTCCTTAC |
| 10_S288C_controls_B6 | ATGTGGTG | TATGCAGT |
| 10_S288C_controls_C1 | CACTATGA | GCGATCTA |
| 10_S288C_controls_C2 | CACTATGA | ATAGAGAG |
| 10_S288C_controls_C3 | CACTATGA | AGAGGATA |
| 10_S288C_controls_C4 | CACTATGA | TCTACTCT |
| 10_S288C_controls_C5 | CACTATGA | CTCCTTAC |
| 10_S288C_controls_C6 | CACTATGA | TATGCAGT |
| 10_S288C_controls_D2 | GCCAGTCA | ATAGAGAG |
| 10_S288C_controls_D3 | GCCAGTCA | AGAGGATA |
| 10_S288C_controls_D4 | GCCAGTCA | TCTACTCT |
| 10_S288C_controls_D5 | GCCAGTCA | CTCCTTAC |
| 10_S288C_controls_D6 | GCCAGTCA | TATGCAGT |

| | | |
|---|---|---|
| 10_S288C_controls_E1 | AATAGCAC | GCGATCTA |
| 10_S288C_controls_E2 | AATAGCAC | ATAGAGAG |
| 10_S288C_controls_E3 | AATAGCAC | AGAGGATA |
| 10_S288C_controls_E4 | AATAGCAC | TCTACTCT |
| 10_S288C_controls_E5 | AATAGCAC | CTCCTTAC |
| 10_S288C_controls_E6 | AATAGCAC | TATGCAGT |
| 10_S288C_controls_F1 | TGCACACA | GCGATCTA |
| 10_S288C_controls_F2 | TGCACACA | ATAGAGAG |
| 10_S288C_controls_F3 | TGCACACA | AGAGGATA |
| 10_S288C_controls_F4 | TGCACACA | TCTACTCT |
| 10_S288C_controls_F5 | TGCACACA | CTCCTTAC |
| 10_S288C_controls_F6 | TGCACACA | TATGCAGT |
| 10_S288C_controls_G1 | CCACCTTG | GCGATCTA |
| 10_S288C_controls_G2 | CCACCTTG | ATAGAGAG |
| 10_S288C_controls_G3 | CCACCTTG | AGAGGATA |
| 10_S288C_controls_G4 | CCACCTTG | TCTACTCT |
| 10_S288C_controls_G5 | CCACCTTG | CTCCTTAC |
| 10_S288C_controls_G6 | CCACCTTG | TATGCAGT |
| 10_S288C_controls_H1 | TGGAATGT | GCGATCTA |
| 10_S288C_controls_H2 | TGGAATGT | ATAGAGAG |
| 10_S288C_controls_H3 | TGGAATGT | AGAGGATA |
| 10_S288C_controls_H4 | TGGAATGT | TCTACTCT |
| 10_S288C_controls_H5 | TGGAATGT | CTCCTTAC |
| 10_S288C_controls_H6 | TGGAATGT | TATGCAGT |
| 11_ETRED_controls_A1 | ATGGAAGC | GGAGTTCC |
| 11_ETRED_controls_A2 | ATGGAAGC | CATGGCCA |
| 11_ETRED_controls_A3 | ATGGAAGC | AATCTCTC |
| 11_ETRED_controls_A4 | ATGGAAGC | TAACCGCG |
| 11_ETRED_controls_A5 | ATGGAAGC | TGGCGGTC |
| 11_ETRED_controls_A6 | ATGGAAGC | CCATCTTA |
| 11_S288C_G8_A7 | ATGGAAGC | ATGTCAAT |
| 11_S288C_G8_A8 | ATGGAAGC | AGTTGGCT |
| 11_S288C_G8_A9 | ATGGAAGC | ACCTAGTA |
| 11_ETRED_controls_B1 | ATGTGGTG | GGAGTTCC |
| 11_ETRED_controls_B2 | ATGTGGTG | CATGGCCA |
| 11_ETRED_controls_B3 | ATGTGGTG | AATCTCTC |
| 11_ETRED_controls_B4 | ATGTGGTG | TAACCGCG |
| 11_ETRED_controls_B5 | ATGTGGTG | TGGCGGTC |
| 11_ETRED_controls_B6 | ATGTGGTG | CCATCTTA |
| 11_S288C_G8_B7 | ATGTGGTG | ATGTCAAT |
| 11_S288C_G8_B8 | ATGTGGTG | AGTTGGCT |
| 11_S288C_G8_B9 | ATGTGGTG | ACCTAGTA |
| 11_ETRED_controls_C1 | CACTATGA | GGAGTTCC |

| | | |
|---|---|---|
| 11_ETRED_controls_C2 | CACTATGA | CATGGCCA |
| 11_ETRED_controls_C3 | CACTATGA | AATCTCTC |
| 11_ETRED_controls_C4 | CACTATGA | TAACCGCG |
| 11_ETRED_controls_C5 | CACTATGA | TGGCGGTC |
| 11_ETRED_controls_C6 | CACTATGA | CCATCTTA |
| 11_S288C_G8_C7 | CACTATGA | ATGTCAAT |
| 11_S288C_G8_C8 | CACTATGA | AGTTGGCT |
| 11_S288C_G8_C9 | CACTATGA | ACCTAGTA |
| 11_ETRED_controls_D1 | GCCAGTCA | GGAGTTCC |
| 11_ETRED_controls_D2 | GCCAGTCA | CATGGCCA |
| 11_ETRED_controls_D3 | GCCAGTCA | AATCTCTC |
| 11_ETRED_controls_D4 | GCCAGTCA | TAACCGCG |
| 11_ETRED_controls_D5 | GCCAGTCA | TGGCGGTC |
| 11_ETRED_controls_D6 | GCCAGTCA | CCATCTTA |
| 11_S288C_G8_D7 | GCCAGTCA | ATGTCAAT |
| 11_S288C_G8_D8 | GCCAGTCA | AGTTGGCT |
| 11_S288C_G8_D9 | GCCAGTCA | ACCTAGTA |
| 11_ETRED_controls_E1 | AATAGCAC | GGAGTTCC |
| 11_ETRED_controls_E2 | AATAGCAC | CATGGCCA |
| 11_ETRED_controls_E3 | AATAGCAC | AATCTCTC |
| 11_ETRED_controls_E4 | AATAGCAC | TAACCGCG |
| 11_ETRED_controls_E5 | AATAGCAC | TGGCGGTC |
| 11_ETRED_controls_E6 | AATAGCAC | CCATCTTA |
| 11_S288C_G8_E7 | AATAGCAC | ATGTCAAT |
| 11_S288C_G8_E8 | AATAGCAC | AGTTGGCT |
| 11_S288C_G8_E9 | AATAGCAC | ACCTAGTA |
| 11_ETRED_controls_F1 | TGCACACA | GGAGTTCC |
| 11_ETRED_controls_F2 | TGCACACA | CATGGCCA |
| 11_ETRED_controls_F3 | TGCACACA | AATCTCTC |
| 11_ETRED_controls_F4 | TGCACACA | TAACCGCG |
| 11_ETRED_controls_F5 | TGCACACA | TGGCGGTC |
| 11_ETRED_controls_F6 | TGCACACA | CCATCTTA |
| 11_S288C_G8_F7 | TGCACACA | ATGTCAAT |
| 11_S288C_G8_F8 | TGCACACA | AGTTGGCT |
| 11_S288C_G8_F9 | TGCACACA | ACCTAGTA |
| 11_ETRED_controls_G1 | CCACCTTG | GGAGTTCC |
| 11_ETRED_controls_G2 | CCACCTTG | CATGGCCA |
| 11_ETRED_controls_G3 | CCACCTTG | AATCTCTC |
| 11_ETRED_controls_G4 | CCACCTTG | TAACCGCG |
| 11_ETRED_controls_G5 | CCACCTTG | TGGCGGTC |
| 11_ETRED_controls_G6 | CCACCTTG | CCATCTTA |
| 11_S288C_G8_G7 | CCACCTTG | ATGTCAAT |
| 11_S288C_G8_G8 | CCACCTTG | AGTTGGCT |

| | | |
|---|---|---|
| 11_S288C_G8_G9 | CCACCTTG | ACCTAGTA |
| 11_ETRED_controls_H1 | TGGAATGT | GGAGTTCC |
| 11_ETRED_controls_H2 | TGGAATGT | CATGGCCA |
| 11_ETRED_controls_H3 | TGGAATGT | AATCTCTC |
| 11_ETRED_controls_H4 | TGGAATGT | TAACCGCG |
| 11_ETRED_controls_H5 | TGGAATGT | TGGCGGTC |
| 11_ETRED_controls_H6 | TGGAATGT | CCATCTTA |
| 11_S288C_G8_H7 | TGGAATGT | ATGTCAAT |
| 11_S288C_G8_H8 | TGGAATGT | AGTTGGCT |
| 11_S288C_G8_H9 | TGGAATGT | ACCTAGTA |
| 12_ETRED_G8_A1 | ATGGAAGC | GAATTCAG |
| 12_ETRED_G8_A2 | ATGGAAGC | CCGGTACG |
| 12_ETRED_G8_A3 | ATGGAAGC | CCGTCATC |
| 12_S288C_G80_A4 | ATGGAAGC | CGTCTATA |
| 12_S288C_G80_A5 | ATGGAAGC | TCAATGAC |
| 12_S288C_G80_A6 | ATGGAAGC | AACGATGC |
| 12_ETRED_G80_A10 | ATGGAAGC | TTGCATGT |
| 12_ETRED_G80_A11 | ATGGAAGC | GGCGCGAT |
| 12_ETRED_G80_A12 | ATGGAAGC | TTAACCGA |
| 12_ETRED_G8_B1 | ATGTGGTG | GAATTCAG |
| 12_ETRED_G8_B2 | ATGTGGTG | CCGGTACG |
| 12_ETRED_G8_B3 | ATGTGGTG | CCGTCATC |
| 12_S288C_G80_B4 | ATGTGGTG | CGTCTATA |
| 12_S288C_G80_B5 | ATGTGGTG | TCAATGAC |
| 12_S288C_G80_B6 | ATGTGGTG | AACGATGC |
| 12_ETRED_G80_B10 | ATGTGGTG | TTGCATGT |
| 12_ETRED_G80_B11 | ATGTGGTG | GGCGCGAT |
| 12_ETRED_G80_B12 | ATGTGGTG | TTAACCGA |
| 12_ETRED_G8_C1 | CACTATGA | GAATTCAG |
| 12_ETRED_G8_C2 | CACTATGA | CCGGTACG |
| 12_ETRED_G8_C3 | CACTATGA | CCGTCATC |
| 12_S288C_G80_C4 | CACTATGA | CGTCTATA |
| 12_S288C_G80_C5 | CACTATGA | TCAATGAC |
| 12_S288C_G80_C6 | CACTATGA | AACGATGC |
| 12_ETRED_G80_C10 | CACTATGA | TTGCATGT |
| 12_ETRED_G80_C11 | CACTATGA | GGCGCGAT |
| 12_ETRED_G80_C12 | CACTATGA | TTAACCGA |
| 12_ETRED_G8_D1 | GCCAGTCA | GAATTCAG |
| 12_ETRED_G8_D2 | GCCAGTCA | CCGGTACG |
| 12_ETRED_G8_D3 | GCCAGTCA | CCGTCATC |
| 12_S288C_G80_D4 | GCCAGTCA | CGTCTATA |
| 12_S288C_G80_D5 | GCCAGTCA | TCAATGAC |
| 12_S288C_G80_D6 | GCCAGTCA | AACGATGC |

| | | |
|---|---|---|
| 12_ETRED_G80_D10 | GCCAGTCA | TTGCATGT |
| 12_ETRED_G80_D11 | GCCAGTCA | GGCGCGAT |
| 12_ETRED_G80_D12 | GCCAGTCA | TTAACCGA |
| 12_ETRED_G8_E1 | AATAGCAC | GAATTCAG |
| 12_ETRED_G8_E2 | AATAGCAC | CCGGTACG |
| 12_ETRED_G8_E3 | AATAGCAC | CCGTCATC |
| 12_S288C_G80_E4 | AATAGCAC | CGTCTATA |
| 12_S288C_G80_E5 | AATAGCAC | TCAATGAC |
| 12_S288C_G80_E6 | AATAGCAC | AACGATGC |
| 12_ETRED_G80_E10 | AATAGCAC | TTGCATGT |
| 12_ETRED_G80_E11 | AATAGCAC | GGCGCGAT |
| 12_ETRED_G80_E12 | AATAGCAC | TTAACCGA |
| 12_ETRED_G8_F1 | TGCACACA | GAATTCAG |
| 12_ETRED_G8_F2 | TGCACACA | CCGGTACG |
| 12_ETRED_G8_F3 | TGCACACA | CCGTCATC |
| 12_S288C_G80_F4 | TGCACACA | CGTCTATA |
| 12_S288C_G80_F5 | TGCACACA | TCAATGAC |
| 12_S288C_G80_F6 | TGCACACA | AACGATGC |
| 12_ETRED_G80_F10 | TGCACACA | TTGCATGT |
| 12_ETRED_G80_F11 | TGCACACA | GGCGCGAT |
| 12_ETRED_G80_F12 | TGCACACA | TTAACCGA |
| 12_ETRED_G8_G1 | CCACCTTG | GAATTCAG |
| 12_ETRED_G8_G2 | CCACCTTG | CCGGTACG |
| 12_ETRED_G8_G3 | CCACCTTG | CCGTCATC |
| 12_S288C_G80_G4 | CCACCTTG | CGTCTATA |
| 12_S288C_G80_G5 | CCACCTTG | TCAATGAC |
| 12_S288C_G80_G6 | CCACCTTG | AACGATGC |
| 12_ETRED_G80_G10 | CCACCTTG | TTGCATGT |
| 12_ETRED_G80_G11 | CCACCTTG | GGCGCGAT |
| 12_ETRED_G80_G12 | CCACCTTG | TTAACCGA |
| 12_ETRED_G8_H1 | TGGAATGT | GAATTCAG |
| 12_ETRED_G8_H2 | TGGAATGT | CCGGTACG |
| 12_ETRED_G8_H3 | TGGAATGT | CCGTCATC |
| 12_S288C_G80_H4 | TGGAATGT | CGTCTATA |
| 12_S288C_G80_H5 | TGGAATGT | TCAATGAC |
| 12_S288C_G80_H6 | TGGAATGT | AACGATGC |
| 12_ETRED_G80_H10 | TGGAATGT | TTGCATGT |
| 12_ETRED_G80_H11 | TGGAATGT | GGCGCGAT |
| 12_ETRED_G80_H12 | TGGAATGT | TTAACCGA |
